# HINS: HLA-mediated immunogenic neoantigen score for robust prediction of immune checkpoint blockade response

**DOI:** 10.64898/2025.12.09.693330

**Authors:** Qiang Yang, Xu Long, Weihe Dong, Xiaokun Li, Kuanquan Wang, Suyu Dong, Gongning Luo, Xianyu Zhang, Tiansong Yang, Xin Gao, Guohua Wang

**Affiliations:** School of Computer Science and Technology, Harbin Institute of Technology, West Dazhi Street, 150001, Harbin, China; Zhengzhou Research Institute, Harbin Institute of Technology, Zhengzhou, 450000, China; School of Computer Science and Technology, Heilongjiang University, Xuefu Road, Harbin 150080, China; Postdoctoral Program of Heilongjiang Hengxun Technology Co., Ltd., Xuefu Road, 150090, Harbin, China; Shandong Hengxun Technology Co., Ltd., Miaoling Road, 266100, Qingdao, China; College of Computer and Control Engineering, Northeast Forestry University, Hexing Road, 150004, Harbin, China; Computer Science Program, Computer, Electrical and Mathematical Sciences and Engineering Division, King Abdullah University of Science and Technology (KAUST), Thuwal 23955-6900, Kingdom of Saudi Arabia; Center of Excellence for Smart Health (KCSH), King Abdullah University of Science and Technology (KAUST), Thuwal, 23955-6900, Kingdom of Saudi Arabia; Center of Excellence on Generative AI, King Abdullah University of Science and Technology (KAUST), Thuwal, 23955-6900, Kingdom of Saudi Arabia; Department of Breast Surgery, Harbin Medical University Cancer Hospital, Haping Road, 150081, Harbin, China; Department of Rehabilitation, the First Affiliated Hospital of Heilongjiang University of Traditional Chinese Medicine, and Traditional Chinese Medicine Informatics Key Laboratory of Heilongjiang Province, Harbin 150040, China

**Keywords:** Immune checkpoint blockade response prediction, Neoantigen quality, HLA evolutionary divergence, Deep attention networks, Anti-tumor immunology, Tumor microenvironment

## Abstract

Immune checkpoint blockade (ICB) therapy response prediction provides remarkable genomic and transcriptomic gains and has successfully understood the molecular mechanisms underlying various cancers. However, only a subset of patients with advanced tumors currently benefits from ICB therapies, which at times incur considerable side effects and costs. Developing predictive tools for ICB response has remained a serious challenge because of the complexity of the immune response and the shortage of large cohorts of ICB-treated patients that include both omics and response data. Here, based on a pooled genomic dataset of 919 patients across multiple solid tumors treated with anti-PD-(L)1 or anti-CTLA-4, we constructed human leukocyte antigen class I (HLA-I)-mediated immunogenic neoantigen score (HINS). HINS is a predictor of ICB response using neoantigen quality and HLA evolutionary divergence to integrate the factors associated with immune activation and evasion. It yielded an overall accuracy of AUC=0.853, outperforming existing predictors and capturing more than 75% true responders. Experimental analysis indicated that patients with higher HINS were more likely to undergo survival benefits following ICB therapy. Our results highlighted the transcriptional and genomic correlations between HINS-identified molecule features and ICB response, such as immunoresponsive gene enrich pathways, favorable and unfavorable genomic subgroups, and hotspot somatic events in driver genes. This study presented an interpretable, accurate deep-learning method using meaningful predictive features to predict the response to ICB, providing biological insights into the complexity of the determinants underlying immunotherapy.

## Introduction

In recent years, immune checkpoint blockade (ICB) therapies have revolutionized the treatment of patients with fatal cancers [1]. These inhibitor agents include antibodies targeting programmed death-1 (anti-PD-1; e.g., nivolumab and pembrolizumab) [2][3] or cytotoxic T lymphocyte-associated antigen 4 (anti-CTLA-4; e.g., ipilimumab) [4], enabling reversal of T regulatory cell-mediated immunosuppression [5]. Durable clinical benefit, however, is limited to a subset of patients. Thus, researchers are searching for predictive biomarkers of ICB therapy response, such as tumor mutational burden (TMB) [6]-[7], T-cell infiltration levels [8], expression of PD-L1 immunohistochemistry [9], somatic genomic features [10], and immune phenotypes [11]. Tumor neoantigen production that escapes T cell central tolerance in the immune system is crucial for antitumor immunity [12]. TMB can partially reflect the situation of neoantigens in patients attributable to genomic-coding mutations. Generally, ICB response-predictive biomarkers are useful but limited, with stratification of patients for immunogenic therapy remaining a hot research topic.

Neoantigens are produced through non-synonymous somatic mutations, creating “non-self” peptides that would be treated as foreign, thus offering immunogenicity to malignant cells [13]-[14]. Tumor neoantigen burden (TNB) is thereby considered an important factor of clinical response to ICB therapy [15]. Researchers are attempting to develop accurate computational tools for the ICB therapy response based on neoantigen immunogenicity, which can activate T-cells to eliminate tumors [16]-[20]. For example, the immuno-predictive score (IMPRS) [17], a predictor of the ICB response in melanoma, was developed on the basis of the immune mechanisms underlying spontaneous regression in neuroblastoma and 15 key immune interactions among immune checkpoint genes. Kim et al. [18] proposed a classifier that can predict ICB inhibitor resistance from functional mutations, using a convolutional neural network (CNN)-based framework to determine the ability of peptides to bind to human leukocyte antigen class I (HLA-I) molecules. However, the relatively high false discovery rate limited the predictive accuracy of these models, as the quality of identified neoantigens is not sufficient. Therefore, Łuksza et al. [19] proposed a neoantigen fitness model of tumor-immune interactions to quantify the immunogenic features of patients’ neoantigen pools under checkpoint blockade immunotherapy. This neoantigen fitness mainly depends on the likelihood of HLA presentation and subsequent T-cell recognition using the relative HLA binding affinity and sequence similarity to identify immunogenic antigens. Inspired by this estimation of neoantigen quality, Han et al. [20] developed the HLA tumor-antigen presentation score (HAPS), by incorporating the HLA-I allele divergence and binding affinity of neoantigens to determine the survival benefit in immunotherapy following ICB therapy. HLA-I allele divergence is a critical element in determining the allele variations within peptide-HLA binding domains. HAPS accessed the immunogenic features of neoantigens using the epitope similarity alignment to the known epitopes from the IEDB website [21]. Nevertheless, the neoantigen immunogenicity of patients calculated by these methods cannot comprehensively reflect the natural traits due to the HLA binding affinity or similarity alignment of epitopes failing to directly explain underlying mechanism of T-cell recognition.

Taking the intricate immune microenvironment into consideration, a multi-modal method could be of significant value for determining ICB therapy efficacy. The quality of tumor-specific neoantigens is primarily determined by their natural characteristics, such as HLA presentation [22], cross-reactivity (a factor that may lead to off-target effects) [23], and T-cell recognition [24]. Therefore, such a comprehensive functional model is needed to satisfy patient stratification of ICB therapy.

Here, we aim to propose a multi-modal method for exploring class I HLA-mediated immunogenic neoantigen score (HINS) by merging HLA-I evolutionary divergence (HED) and neoantigen quality. HINS uses deep attention networks (DANs) to calculate the ‘non-self’ recognition potential of a neoantigen and combines the self-discrimination of immune system (i.e., HLA presentation, T cell cross-reactivity). HINS-identified tumor microenvironment (TME) features are also analyzed. Furthermore, we perform comprehensive investigations to identify genomic biomarkers for co-evolutionary assessments of ICB response and resistance. Our findings demonstrated that HINS can be a valuable tool for predicting the immune response to ICB therapy.

## Materials and methods

### Cohorts for ICB therapy response

In this study, we constructed a large-scale patient cohort for predicting the response to ICB therapy (anti-PD-(L)1, anti-CTLA-4, or a combination of these therapies). In total, 919 patients with whole-exome sequencing (WES) data were used to train and validate the developed predictor (**Figure 1A**, **Table 1**). Patients’ samples were collected from eight previously published cohorts [20], [25]-[31] across six cancer types [skin cutaneous melanoma (SKCM), n=359; non-small cell lung cancer (NSCLC), n=519; head and neck squamous cell carcinoma (HNSCC), n=12; bladder cancer, n=27; anal cancer, n=1; sarcoma, n=1]. The clinical characteristics of the constructed cohorts were presented in **Table S1**. These studies were conducted in accordance with the Declaration of Helsinki. All enrolled patients’ information was public and approved by the ethics committees of the National Cancer Center. **Table 1** displayed the statistical traits of patient cohorts.

**Figure 1.**
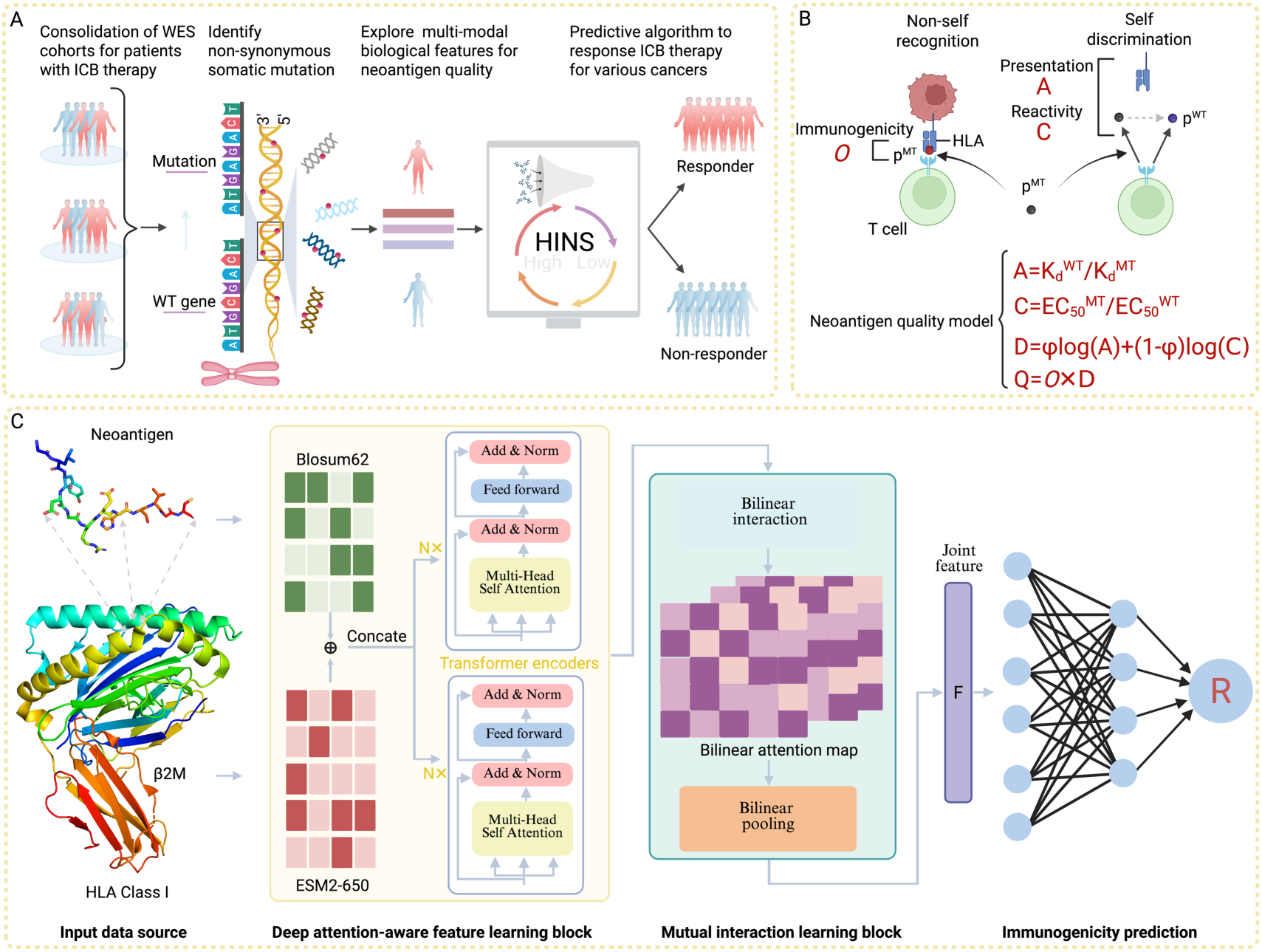
Rationale and definition of HLA-I-mediated immunogenic neoantigen score (HINS). **A**. Overview of the HINS method for immune checkpoint blockade (ICB) response prediction. We pooled 9 cohorts of ICB recipients with corresponding whole-exome sequencing (WES) and Response Evaluation Criteria in Solid Tumors (RECIST) classification. The neoantigen quality and HLA divergence were integrated to support the model inference. **B**. The visualized workflow of the neoantigen quality model, including neoantigen immunogenicity, HLA presentation and T cell cross-reactivity. **C**. The structure of the deep attention networks is applied to learn the immunogenicity of HLA-I-bound neoantigen, which plays a central role in host immune response.

**Table 1.** Characteristics of cohorts utilized in this study.

| Cohorts | Cancers | Samples,<br>n(%) | Responders | Non-responders | Sources |
| --- | --- | --- | --- | --- | --- |
| Miao1 | NSCLC&<br>SKCM | 208(22.7%) | 57 | 151 | [25] |
| Miao2 | Other | 41(4.5%) | 17 | 24 | [25] |
| Anagnostou | NSCLC | 89(9.7%) | 41 | 48 | [26] |
| Rizvi | NSCLC | 34(3.6%) | 12 | 22 | [27] |
| Ravi | NSCLC | 309(33.5%) | 121 | 188 | [28] |
| Wang | NSCLC | 30(3.3%) | 7 | 23 | [20] |
| Roh | SKCM | 53(5.7%) | 8 | 45 | [29] |
| Van | SKCM | 100(11%) | 22 | 78 | [30] |
| Synder | SKCM | 55(6%) | 13 | 42 | [31] |

We evaluated clinical responses using RECIST (Response Evaluation Criteria in Solid Tumors) version 1.1 [32]. Patients treated with ICB therapy were categorized as responder (response with durable clinical benefit (DCB)) and non-responder (response with non-clinical benefit (NCB), **Figure S1A-C**). DCB was defined as complete response (CR) or partial response (PR). NCB was defined as progressive disease (PD) or stable disease (SD). All clinical data, such as overall survival (OS), progression-free survival (PFS), and clinical response, were obtained from previous studies. OS was defined as the time from the beginning of ICB therapy to death from any cause. PFS was defined as the time from the beginning of ICB therapy to the date of cancer progression or death, whichever is earlier. Tumors from patients with PFS or OS of less than 4 weeks were excluded from the analysis, as these patients might have had a disease that was too advanced to experience a clinical benefit from ICB. Additionally, we stratified the analysis of the distribution of TMB across the eligible tumor types (**Figure S1D**).

### Whole exome sequencing and analysis

For cohorts with WES data (**Figure S2**), we aligned the raw fasta files with reference human genome GRCh38 using Burrows-Wheeler Aligner (BWA, http://bio-bwa.sourceforge. net) version 0.7.17 [33]. Picard (v 2.26.6) was employed to remove duplicate reads. Base quality score recalibration and read realignment were performed using the Genome Analysis Toolkit (GATK, https://software.broadinstitute.org/gatk) version 4.1 [34]. Somatic single nucleotide variants (SNVs) were identified using MuTect2 [35]. Candidate tumor mutations were selected only when the following criteria were met: 1) detected in high-quality reads more than 5 times; 2) with a variant allele frequency (VAF) higher than 1%; 3) not observed in several single-nucleotide polymorphism databases (e.g., 1000 Genomes Project with version phase 3, dbSNP version 137 and COSMIC database). Somatic mutations were detected using GATK4 pipelines in the Terra cloud platform (https://app.terra.bio/).

### Neoantigen immunogenicity prediction

The immunogenic potential of the non-synonymous somatic mutations was assessed by calculating the immunogenicity between HLA-I alleles and mutant epitopes. According to our previous studies [36]-[37], we constructed DANs to predict HLA-I-restricted neoantigen immunogenicity (**Figure 1C**), which implies T-cell recognition.

#### Input embedding

The embedding matrix Blosum62 was employed to convert the one-dimensional (1D) amino acid sequences for the peptides into 2D matrices. Let ℓ be the maximum length of input peptides and 𝜐 be the embedding dimension of amino acids, respectively. Peptides shorter than ℓ were padded with zeros. For the HLA-I molecular pseudosequence, we carefully selected 35 amino acid residues that are most likely to interact with peptides to encode HLA sequences. Hence, the input embedding of a given pair of peptide 𝑃*_B_* ∈ ℝ^ℓ×𝜐^ and HLA molecular pseudosequence 𝑄*_B_* ∈ ℝ^35×𝜐^ can be formulated as follows:

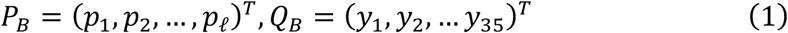

where 𝑝*_i_* ∈ ℝ^𝜐^ indicates the embedding representation of the 𝑖*_th_*, amino acid of a peptide and 𝑞*_i_* ∈ ℝ^𝜐^ is the embedding representation of the 𝑗*_th_* amino acid of allele pseudosequence.

Additionally, we utilized the pretrained protein language model ESM2 with 650 million parameters to generate embeddings of peptide and HLA as complementary domain knowledge [38]. ESM2 outputted embeddings with a length 1280 for individual amino acids in a sequence. Embeddings of peptide 𝑃*_E_* ∈ ℝ^ℓ×1280^ and HLA molecular pseudosequence 𝑄*_E_* ∈ ℝ^35×1280^ were obtained. Blosum62 and ESM2 embeddings were concatenated to fully indicate the high-level features in distinct biological significances, thus, representations 𝑃 ∈ ℝ^ℓ×(𝜐+1280)^ and 𝑄 ∈ ℝ^35×(𝜐+1280)^ were obtained.

#### Deep attention-aware feature learning block

Recently, attention-based deep learning models have remarkably succeeded in computer vision, natural language processing and especially in computational biology [36]-[37], [39]. We used deep attention-aware extractors to separately learn the biological and sequential features of short peptides and HLA pseudosequence. Specifically, after the concatenated embedding for a given peptide, we fed it into three consecutive molecule transformer encoders (MTEs), which can be described as follows:

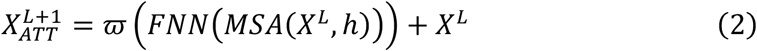

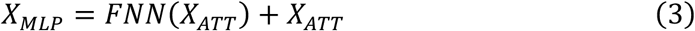

where 𝑋*_ATT_* and 𝑋*_MLP_* denote the output features of multi-head attention and fully connected network (FNN) layer in the MTE, respectively. 𝜛(·) is the activation function 𝑅𝑒𝐿𝑢. 𝑀𝑆𝐴(·) indicates the multi-head attention function and $h$ is the corresponding head number. After 𝑀𝐴𝑆(·) or 𝐹𝑁𝑁(·) function, there are the following residual layer and layer normalization in each transformer encoder. Let 𝐿 be the number of the MTE and when 𝐿 is equal to 0, 𝑋*^L^* degrades into 𝑃. The high-level biological features of peptides 𝑋*_pep_* ∈ ℝ^ℓ×*t*^, where 𝑡 denotes the feature dimension, are finally represented by a multilayer perceptron following the consecutive MTEs. Similarly, the superior features of the HLA-I molecule 𝑋*_HLA_* ∈ ℝ^35×*t*^ is also obtained.

#### Mutual interaction learning block

Bilinear attention networks were introduced to extract mutual local interactions between HLA-I molecules and short peptides. Specifically, we first built a bilinear interaction map using the biological features 𝑋*_pep_* and 𝑋*_HLA_* to construct a single head mutual interaction matrix 𝑀 ∈ ℝ^ℓ×35^ as follows:

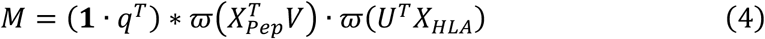

where 𝑉 ∈ ℝ^ℓ×*K*^ and 𝑈 ∈ ℝ^35×*K*^ are learnable weight matrices for peptide and HLA pseudosequence representations, 𝟏 ∈ ℝ^ℓ^ and 𝑞 ∈ ℝ*^K^* are a fixed vector of ones and learnable weight vector, respectively, and ∗ denotes element-wise product.

Subsequently, we introduced a bilinear pooling layer to calculate the joint representation 𝑟′ ∈ ℝ*^K^* as follows:

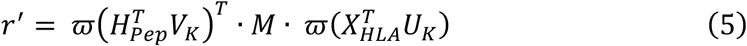

where 𝑉*_K_* and 𝑈*_K_* indicate the 𝑘*_th_* column of weight matrices 𝑉 and 𝑈, respectively. Of note, there are no additional learnable parameters in this pooling layer. Furthermore, a sum pooling on the joint representation 𝑟′ was applied to obtain a compact feature map as follows:

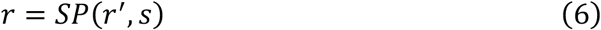

where the 𝑆𝑃(·) function represents a non-overlapped and 1D sum pooling algorithm with stride 𝑠. It decreased the dimensionality of 𝑟′ ∈ ℝ*^K^* to 𝑟 ∈ ℝ*^K^*^/*s*^.

#### Immunogenicity prediction

To calculate the immunogenicity probability, we fed the obtained joint representation 𝑟 into a tri-layer fully connected network and a sigmoid function.

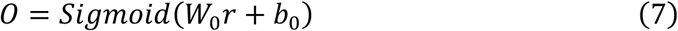

where 𝑊_0_ and 𝑏_0_ are learnable weight matrix and bias vector, respectively. We minimized the distance between the predicted immunogenicity and the ground-truth using the binary cross-entropy loss function as follows:

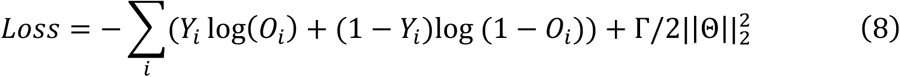

where Θ indicates the set controlling all the learnable weight and bias parameters mentioned above, 𝑌*_i_* is the real label of the 𝑖*_th_* input samples, and 𝑂*_i_* is the final predicted score. Γ stands for a hyperparameter for L2 regularization.

### Multi-modal features for HINS construction

According to previous studies [40], HLA-I divergences with loss of heterozygosity (LOH) may affected tumor-specific neoantigen presentation. Thus, by including the divergence in paired HLA-I alleles [41], we introduced a score for assessing response to immunotherapy. The HLA-I-mediated immunogenic neoantigen score (HINS) was defined as follows:

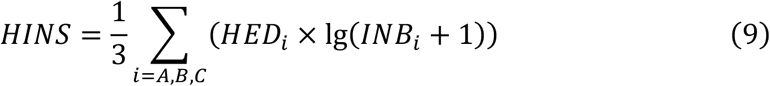

where 𝐻𝐸𝐷*_i_* indicates the HLA divergences of alleles A, B and C, which is calculated by the Grantham distance metric [42]. 𝐼𝑁𝐵*_i_* denotes the immunogenic neoantigen burden for each HLA allele. We defined *INB* as the number of neoantigens with high quality (𝑄 = 𝑂 × 𝐷), where 𝑂 is the neoantigen immunogenicity estimated by the constructed deep attention-wise model and 𝐷 is the self-discrimination to model the immune system properties (i.e., HLA presentation and TCR cross-reactivity). Self-discrimination is quite important yet always neglected by most studies, which can overcome the principles of negative T-cell selection. With this conception, we built self-discrimination 𝐷 between a mutant neopeptide (𝑝*^MT^*) and its corresponding wild-type peptide (𝑝*^WT^*) as follows:

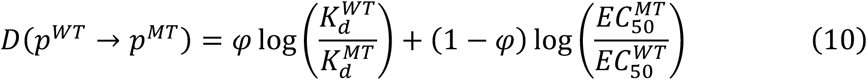

where 𝜑 is a relative weight between the terms of presentation and cross-reactivity. 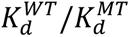 represents differential HLA presentation. 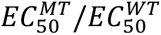 is the cross-reactivity distance. Therefore, by integrating multi-model features, i.e., HLA divergences, neoantigen immunogenicity, HLA presentation and TCR cross-reactivity, HINS was developed to predict and analyze the ICB response in patients with cancer.

### Training implementation

For immunogenicity prediction model training, the used hyperparameters including learning rate [1e-4, 1e-3, 1e-2, 0.05, 0.1], batch size [32, 64, 128, 256], epoch [10, 30, 50, 100], L2 regularization [0.0001, 0.001, 0.01] and dropout [0.1-0.9]. The learnable model parameters were calculated using standard backpropagation. Stochastic gradient descent was employed to replace Adam to optimize the objective function because Adam was unstable in our training process. This model was trained on Python 3.10 and Pytorch 2.2.1 on CUDA 12.1 with NVIDIA RTX 8000 GPU (64 GB of RAM). Of note, the graphic memory of the computing device must be at least 48 GB for the large protein language model ESM2. More detailed information on model development is presented in the **Table S2**.

## Results

### Accurate prediction of neoantigen immunogenicity

Neoantigen immunogenicity has been proven to be a fundamental factor in the immunogenic response of patients who received ICB therapy. In this study, we developed a deep attention networks (DAN) model to predict the immunogenicity of class I HLA-bound neoantigens. Training data were collected from experimentally verified samples from the IEDB website [21], which recorded 66,298 neoantigen immunogenicity hits (positive=49,217 samples, negative=17,081 samples, **Table S3**). To balance the ratio of positive/negative ratio, 10-fold length-matched decoys were randomly generated from the human proteome as negative examples. We employed the 5-fold cross-validation in the training process and chose the model with the best results to evaluate the predictive performance. The validation sets were constructed using 5 collected ICB cohorts [26]-[31] with two types of tumors, i.e., SKCM and NSCLC. More than 148,000 neoantigen immunogenicity samples were obtained (**Table S4**). Of note, we employed high-affinity binding as a proxy for the immunogenicity ground-truth (IC50 < 50nM). To comprehensively evaluate the predictive performance of the proposed DAN model, four golden standard metrics were utilized, i.e., the area under the receiver operating characteristic curve (AUC), the area under the precision-recall curve (AUPR), the positive prediction value at top-n (PPVn), and the F1-score (**File S1**).

We compared DAN with three state-of-the-art (SoTA) predictive tools for the immunogenicity of class I restricted antigens, i.e., NetMHCpan-4.1(represented as Netpan in the following text) [43], CNN [18] and HLAthena [44] (**File S1**). The evaluated average results and standard deviation of SKCM and NSCLC neoantigen immunogenicity prediction are presented in **Table 2**. As shown, the proposed model has consistently outperformed other advanced tools in all four metrics, especially in AUPR, which is more rigorous and reliable for unbalanced data training. For instance, in the neoantigen immunogenicity prediction of SKCM cancer, DAN yielded 11.2%, 17.5%, 21.7%, and 14.7% performance gains in terms of the utilized metrics, respectively, against the runner-up tool CNN (𝑃 < 0.05, with statistical significance). To further explore the representational gains, we conducted an ablation study to assess their unique and combined contributions by constructing three variants of DAN (BLOSUM62, ESM2 and the combination). The results showed that the concatenated embedding consistently yielded the best performance across all datasets (**Table S5**), indicating the combination variant provides both conserved biochemical substitution patterns and contextual immunogenicity-related feature representations, thereby enhancing immunogenicity prediction. The high improvement is mainly attributed to our well-designed attention-aware representation, which can effectively capture the latent features of each amino acid residue of HLA-I neoantigens, indicating an intrinsic immunogenicity mechanism.

**Table 2.** Comparison of four predictive tools among SKCM and NSCLC cancer regarding AUC, AUPR, PPVn and F1-score values.

|  | Method | AUC | AUPR | PPV | F1-score |
| --- | --- | --- | --- | --- | --- |
| SKCM | Netpan | 0.694(0.013) | 0.196(0.043) | 0.217(0.031) | 0.329(0.029) |
|  | CNN | 0.785(0.009) | 0.372(0.025) | 0.437(0.018) | 0.552(0.013) |
|  | HLAthena | 0.736(0.007) | 0.344(0.017) | 0.412(0.014) | 0.539(0.017) |
|  | DAN | **0.873(0.003)** | **0.437(0.009)** | **0.532(0.007)** | **0.633(0.005)** |
| NSCLC | Netpan | 0.675(0.012) | 0.257(0.047) | 0.239(0.027) | 0.331(0.026) |
|  | CNN | 0.782(0.010) | 0.356(0.022) | 0.431(0.015) | 0.532(0.011) |
|  | HLAthena | 0.717(0.005) | 0.312(0.023) | 0.442(0.015) | 0.454(0.013) |
|  | DAN | **0.842(0.002)** | **0.425(0.011)** | **0.513(0.006)** | **0.627(0.004)** |

*Note:* Best result is bolded.

Generally, the simple data-splitting strategy in the training process might leak prior knowledge for the predictive learners, and the tested results are thus usually overestimated. For this reason, we independently utilized Miao et al’s neoantigen data as training data to train the predictors and tested them on the five previous cohorts and Miao et al.’s cohort (**Table S6**). Of note, the neoantigens in Miao’s dataset were from 269 patient tumors across 6 cancer types. Only the neoantigens from SKCM and NSCLC were used as the training samples, whereas the neoantigens produced by other cancers (HNSCC, bladder cancer, anal cancer, and sarcoma) were used to evaluate the predictive performance. Consequently, DAN still exhibited competitive superiority versus the compared prediction tools. Intuitively, our model accurately identified the immunogenicity of neoantigens in more than 90.2% of the test candidates in each cohort of common cancers (**Figure S3A-B**). Meanwhile, DAN manifested strong generalization capability in predicting the neoantigen immunogenicity for rare cancers that have never occurred in the inference stage (**Figure S3C**). A two-sided Wilcoxon test was conducted for this evaluation, and the P-value was less than 0.001, indicating the comparisons were statistically significant. These results demonstrated that the tailored deep-learning model could accurately identify the immunogenicity of neoantigens and be a fundamental pillar to support the HINS construction.

### HINS improves immunotherapy response prediction

To evaluate whether the HINS identifies clinical outcomes of treated patients, we utilized Miao1 and Ravi et al.’s cohorts as training set. We assigned a continuous cutoff point of 14.5 for high vs. low HINS, which was stratified according to the maximum Youden’s index ( 𝐽 = 𝑠𝑒𝑛𝑠𝑖𝑡𝑖𝑣𝑖𝑡𝑦 + 𝑠𝑝𝑒𝑐𝑖𝑓𝑖𝑐𝑖𝑡𝑦 − 1) [45] in the pooled training cohorts. The threshold was subsequently applied unchanged to all independent validation cohorts. Previous studies demonstrated that the predictive tools built on biological scores [46]-[47], such as cytolytic factor [47], generalized better than the machine learning-based predictors constructed on transcriptomic signatures. We first compared the predictive performance of HINS with that of current transcriptome-based and machine learning-based predictors. These predictors of response to immunotherapy for transcriptome-based predictors were generated using Auslander et al.’s method [17]. The resulting performance indicated that HINS (AUC=0.853) was superior to these tools, with 6.2% overall improvement against the runner-up method HAPS (**Figure 2A**). Apparently, custom-made deep-learning predictor is competitive to transcriptome-based and machine learning-based ones, consistent with Kim et al.’s research [18]. The number of false and true positives (responders) and false and true negatives (non-responders) at different HINS binary thresholds showed that our method can capture approximately 75% of true responders, greatly alleviating the false discovery rate compared with the findings in previous studies (**Figure 2B**).

**Figure 2.**
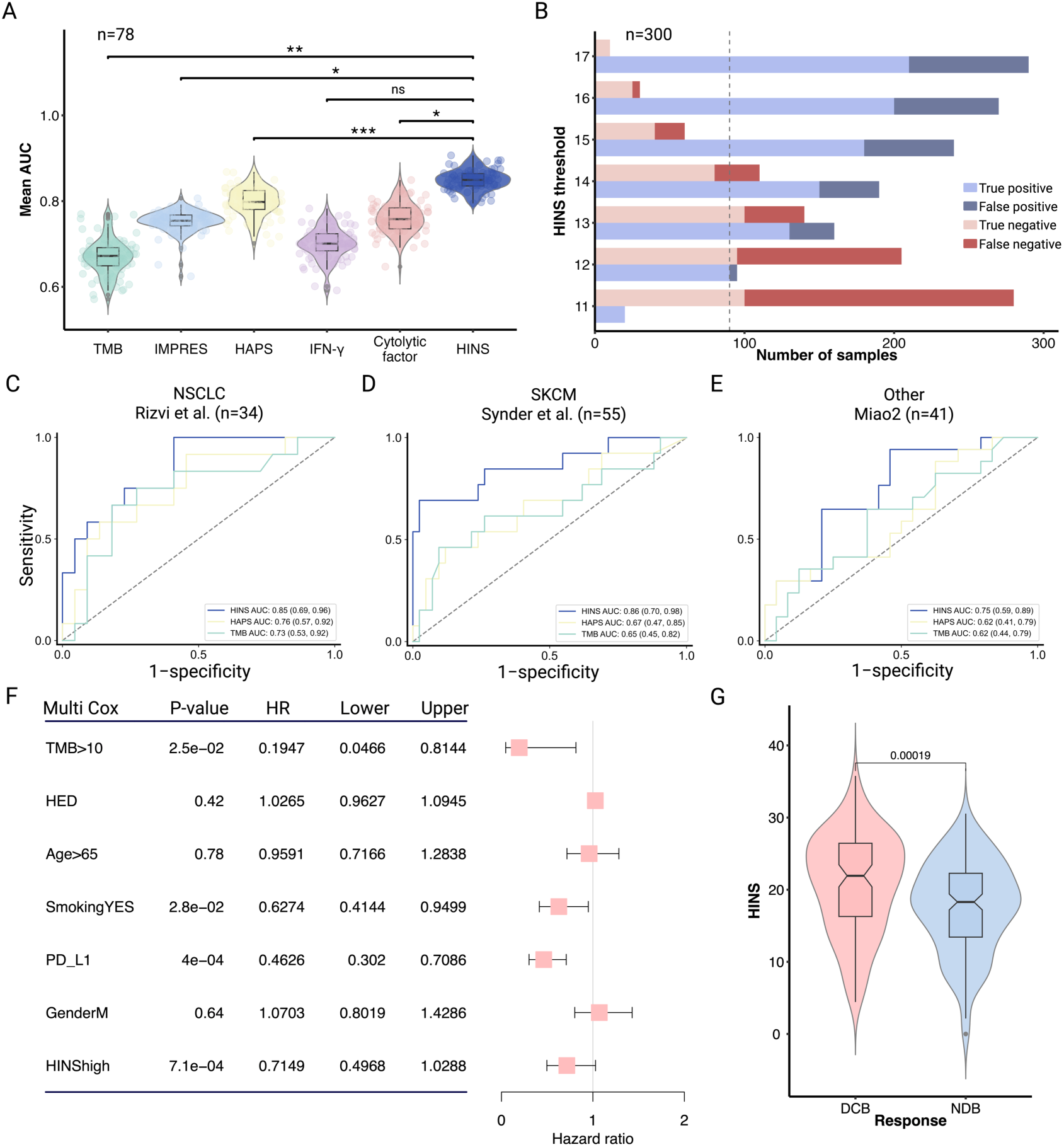
HINS significantly improves the predictive performance of the response to immunotherapy. **A**. AUC results of HINS and traditional published predictors across five publicly available ICB treatment datasets. **B**. Bar plots indicate the prediction accuracy and error types for different HINS thresholds (where a positive label corresponds to a ‘responder’ prediction) on the aggregate compendium of the 300 participants in seven datasets. **C-E**. ROC curves and corresponding AUCs with 95% concordance index (CI) of HINS (blue curves), HAPS (yellow curves) and the TMB biomarker (green curves) on the training sets and across various unseen validation sets. **F**. Multivariate Cox analysis showing significantly prolonged progression-free survival in the high HINS subgroup (n = 772). The error bars indicate 95%CI for HR. **G**. Significant difference in HINS between durable clinical benefit and non-durable benefit subgroups using the two-sided Wilcoxon test (n=919).

We then investigated whether above observation also held true when comparing the performance of published ICB cohorts among three distinct cancer types, including SKCM (Rizvi et al., Wang et al., and Anagnostou et al.), NSCLC (Synder et al., Roh et al., and Van et al.) and other cancers (Miao2). The resulting AUCs ranged 0.43-0.88 (**Figure 2C-D**, **Figure S4**). HINS consistently surpassed HAPS and TMB biomarker almost across all cohorts, indicating that our method is more accurate in assessing patient survival (two-tailed DeLong’s test, 𝑃 < 2 × 10^-4^). Even for cancer types not observed in the training process, such as HNSCC and bladder cancer in the ‘other cohort’, HINS still exhibited significantly high performance (**Figure 2E**).

To estimate whether HINS was associated with clinical benefit, we first performed univariate Cox regression in an independent cohort of 772 ICB-treated patients with multi-parameter information (**Table S7**). In this model, higher HINS scores were significantly associated with prolonged progression-free survival ( 𝑃 = 1 × 10^-4^; 𝐻𝑅 = 0.70, 95%𝐶𝐼 = 0.59 − 0.84; **Figure S5A**), and a similar beneficial trend was observed for overall survival(**Figure S5B-C**). We then constructed a multivariate Cox proportional hazards model to determine whether HINS served as an independent prognostic factor. The model was adjusted for TMB>10 (>10 vs. <10 mutants/exome), HED, Age>65 (>65 vs. < 65), SmokingYES (Yes vs. Not), PD-L1, GenderM (Male vs. Female), and HINShigh (high vs. low). As observed, higher HINS ( 𝑃 = 7.1 × 10^-4^; 𝐻𝑅 = 0.71, 95%𝐶𝐼 = 0.50 − 1.03) were more prone to associate with improved progression-free survival (PFS, **Figure 2F**). The biological factor PD-L1 exhibited a clearer association with statistical significance ( 𝑃 = 4 × 10^-4^; 𝐻𝑅 = 0.46, 95%𝐶𝐼 = 0.3 − 0.71). However, the evaluation of PD-L1 was constrained by sparse data availability, restricting its assessment to a small patient subset. Conversely, HINS was uniformly derived across the entire WES landscape. Thus, we position HINS as a broadly applicable predictor. Furthermore, a positive association was illustrated between the HINS and durable clinical benefit (DCB) after ICB therapy by exploring the predictive value for ICB response of HINS in most tumor types, e.g., SKCM and NSCLC (𝑃 = 1.9 × 10^-4^, DCB vs. NCB: 21.93 vs. 18.3; **Figure 2G**). In summary, HINS provided monotonic prediction of response to immunotherapy and displayed positive associations with efficacy and clinical benefits.

### HINS facilitated better patient stratification

We further surveyed whether HINS could stratify the prediction of immunotherapy outcomes of patients. First, our Kaplan-Meier (K-M) analysis revealed that high HINS (cutoff with 14.5) had significantly better survival compared to those with low scores in almost all cohorts. Even though our cutoff was determined only from the training set, supporting its robustness as a global operating point across cancer types. Taken common cancers for example, the OS outcome is greatly improved in NSCLC (Wang et al.: 𝑃 = 0.047, 𝐻𝑅 = 0.45, 95%𝐶𝐼 = 0.2 − 1.03; Rizvi et al.: 𝑃 = 0.016, 𝐻𝑅 = 0.13, 95% = 0.05 − 0.34) and SKCM (Roh et al.: 𝑃 = 0.030, 𝐻𝑅 = 0.5, 95%𝐶𝐼 = 0.23 − 1.09; Van et al.: 𝑃 = 0.13, 𝐻𝑅 = 0.67, 95%𝐶𝐼 = 0.42 − 1.08; **Figure 3A-B, Figure S6A-F**). For the unseen cancers, our method showed a powerful generalization ability (Miao2: 𝑃 = 0.048, 𝐻𝑅 = 0.34, 95%𝐶𝐼 = 0.14 − 0.87, **Figure 3C**). Although HINS in these independent cohorts exhibited promising trends consistent with the general pattern, limited sample sizes might constrain statistical power. To rigorously validate reliability, we performed a pooled analysis by integrating independent validation cohorts within specific cancer types. In the aggregate NSCLC validation set (Rizvi and Wang cohorts, n=64), high HINS remained strongly associated with prolonged OS (𝑃 = 0.015, 𝐻𝑅 = 0.46, 95%𝐶𝐼 = 0.25 − 0.84; **Figure 3D**). Similarly, the pooled SKCM validation set (Roh, Van and Synder cohorts, n=208) confirmed the robustness of HINS in stratifying survival outcomes ( 𝑃 = 0.049, 𝐻𝑅 = 0.71, 95%𝐶𝐼 = 0.51 − 1; **Figure 3E**). However, HINS failed to infer the OS survival result in Anagnostou et al.’s cohort (𝑃 = 0.821, 𝐻𝑅 = 0.93, 95%𝐶𝐼 = 0.48 − 1.78; far from statistical significance, **Figure S6F**) due to neoantigen quality was hard to learn in a patient with low tumor purity. Specifically, low tumor purity significantly limits the sensitivity of somatic mutation calling from WES data, resulting in an incomplete retrieval of the neoantigen landscape. Since HINS is constructed from high-quality neoantigen and HED features, such the omission of neoepitopes in low-purity samples inevitably compromises the model’s accuracy in stratifying responders. The PFS results were also recorded (**Figure S7**). While HINS was designed as a multiple cancer predictor, we also explored the potential of cancer-specific optimization for generalizability conformation. We constructed an NSCLC-specific model (cutoff = 18.5) that trained on the large-scale NSCLC cohort Ravi et al., n=309). The NSCLC-specific HINS model yielded consistent predictive performance and significant survival stratification (**Figure S8**), further confirming that the neoantigen quality features captured by HINS are robust drivers of immunogenicity within specific cancer types.

**Figure 3.**
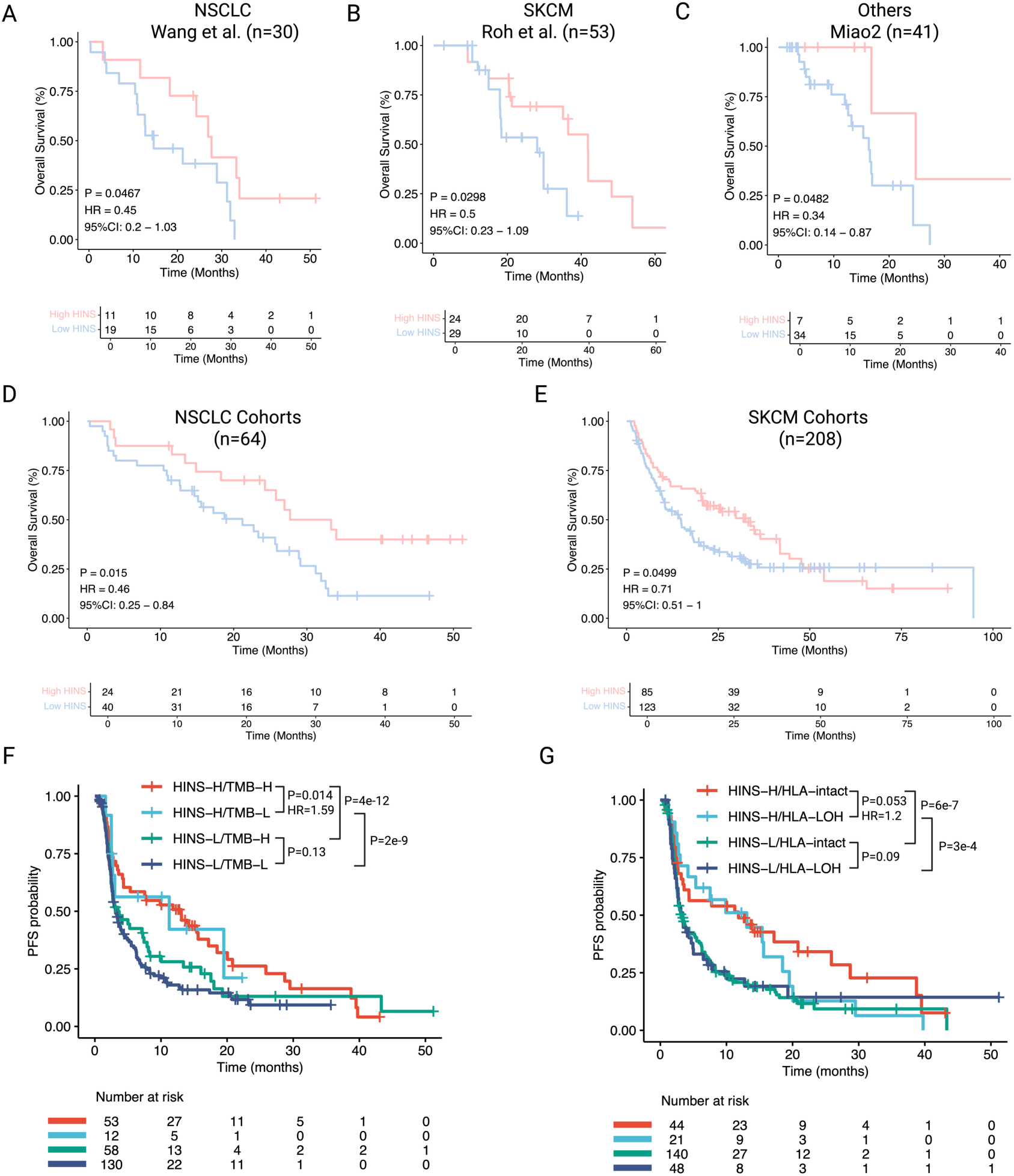
HINS predicts patient outcomes following ICB therapy with powerful stratification. **A-C**. Kaplan-Meier analysis indicates significant correlations between high HINS and improved overall survival after immunotherapy in multiple cancer types. Wang et al.’s NSCLC cohort (A), Roh et al.’s SKCM cohort (B), and Miao et al.’s cohort including multiple cancers (C). **D-E**. Validation in pooled datasets to enhance statistical robustness. K-M analysis of the pooled NSCLC cohorts (n=64) (D) and pooled SKCM cohorts (n=208) (E) confirms that high HINS remains a significant predictor of prolonged OS in larger, integrated populations. **F-G**. Kaplan-Meier analysis of PFS when combining HINS and TMB (F) or HLA-LOH (G). The P-value was determined by univariable Cox proportional hazards regression. H: high, L: low.

To investigate the efficiency of the combined use of HINS and traditional biomarkers, such as TMB and LOH at the HLA-I locus (HLA-LOH), we calculated HRs for them (**Table S8**). Regarding TMB, our results showed that participants with concurrently high HINS and high TMB had the longest PFS (𝑃 = 0.014, 𝐻𝑅 = 1.59, **Figure 3F**). Of note, HINS identified substantial participants with low-TMB that could respond to ICB therapy at a similar benefit to participants with high-TMB. For HLA-LOH, an important factor associated with immunotherapy resistance and immune escape, we studied the prevalence of available HLA-LOH data. Our K-M analysis revealed that the participant group with high HINS and HLA-LOH had significantly worse survival compared to those with high HINS and HLA-intact (no LOH happened at the HLA-I locus, 𝑃 = 0.053, 𝐻𝑅 = 1.20, **Figure 3G**). Similar results were observed for OS (**Figure S6G-H**). Evidently, patients with high HINS, regarding whether they showed high or low levels of TMB and HLA-LOH, tended to obtain homologous benefits from immunotherapy. Taken together, HINS allowed for better patient stratification in terms of predicted ICB outcomes, especially for the combination of HINS and other important biological factors.

### HINS correlates with T cell immune response

To further verify the biological relevance of HINS, we analyzed the T-cell receptor (TCR) repertoire using WES data from previous published work (n=81) [20]. We observed a significant positive correlation between HINS and both TCR diversity (Spearman 𝑅 = 0.266, 𝑃 = 0.017 ; **Figure 4A**) and clonality (Spearman 𝑅 = 0.282, 𝑃 = 3.09 × 10^-5^ ; **Figure 4B**, **Table S9**), indicating that HINS-predicted neoantigens effectively drive T-cell clonal expansion. Beyond clonal abundance, we assessed the similarity of the immune repertoire via TCR edit (Hanming) distance. Patients in the high-HINS subgroup exhibited significantly higher edit distances compared to the low-HINS subgroup (𝑃 < 2.26 × 10^-9^; **Figure 4C**, **Table S10**). This similarity alignment remained significant when stratifying patients into four subgroups based on combined HINS and diversity status (𝑃 < 0.009; **Figure 4D**). This result suggested that high-quality neoantigens foster a more diverse TCR repertoire capable of recognizing a broader spectrum of epitopes.

**Figure 4.**
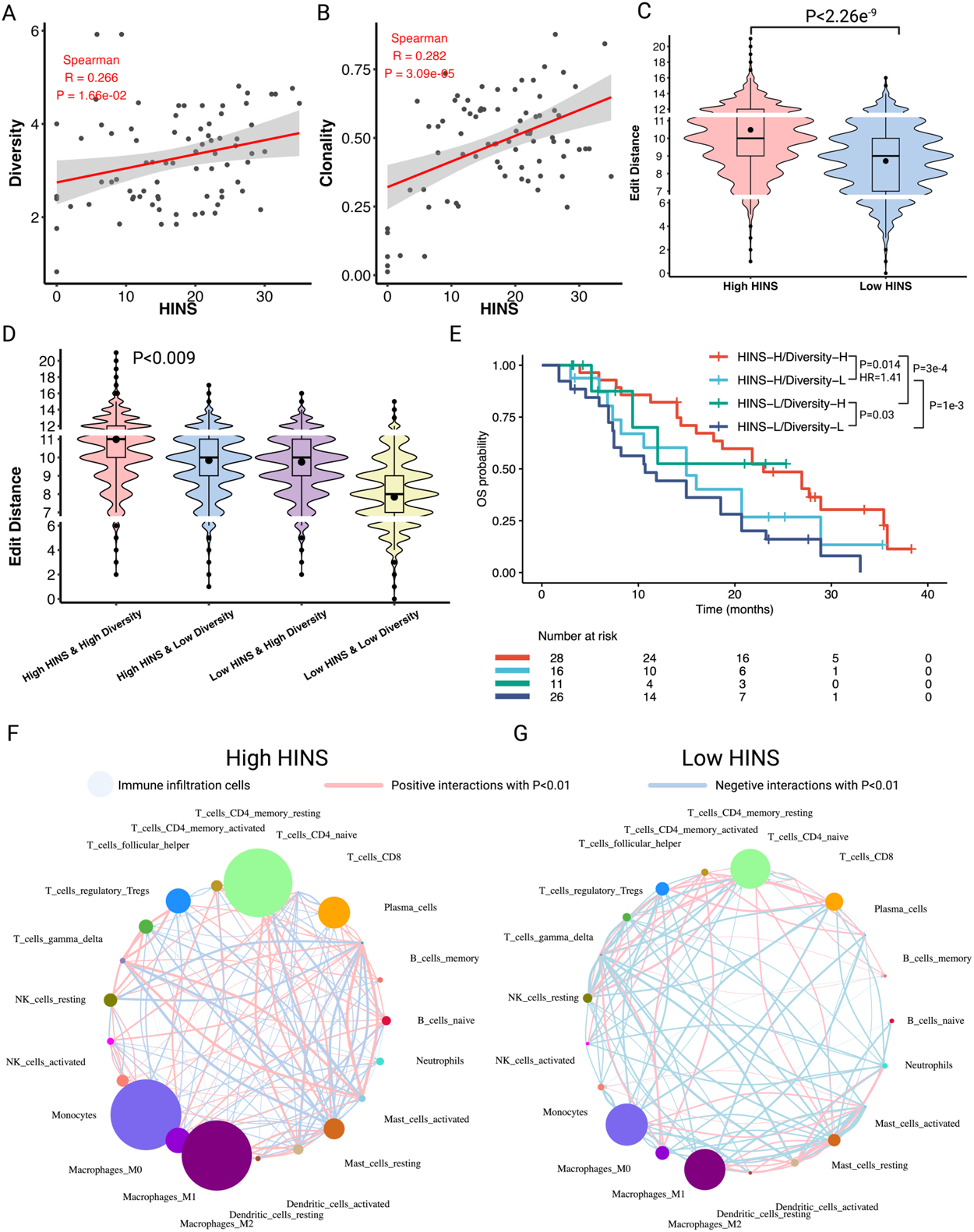
HINS is correlated with actual T cell immune response. **A-B**. Positive correlation was found between HINS and TCR diversity (A) and TCR clonality (B). The red line represents the linear fit with a 95% confidence interval (shaded area). Spearman correlation coefficients (R) and P-values are indicated. **C**. Violin plots displaying the distribution of TCR edit distances in high (red) versus low (blue) HINS subgroups. **D**. Distribution of TCR edit distances across four subgroups stratified by HINS and TCR diversity. In the box plots, the dots represent mean values, the center line corresponds to median, box boundaries correspond to the first and third quartiles, the upper whisker is max and lower whisker is min. **E**. Kaplan-Meier survival analysis of OS for patients stratified by the combination of HINS and TCR diversity status (High vs. Low). The High HINS & High Diversity group (red curve) showed superior survival benefits. **F-G**. Interactions between immune cell types within the tumor microenvironment. The size of the point indicates the level of immune infiltration. The thickness of the line indicates the strength of the associations evaluated via Spearman correlation.

More important, the combination of HINS and TCR diversity significantly improved prognostic stratification (**Figure 4E**). Patients with concurrently high HINS and high TCR diversity yielded the most favorable OS, significantly outperforming the high HINS and low TCR diversity group (𝑃 = 0.014, 𝐻𝑅 = 1.41), or low HINS and high TCR diversity group (𝑃 = 3 × 10^-4^). These observations confirm that HINS learns a distinct dimension of immunogenicity that, when combined with TCR immune response, offers a more holistic and accurate prediction of ICB efficacy.

### HINS correlates with immune transcriptional programs

We next focused on the relationship between HINS and the immune transcriptional programs in the tumor microenvironment (TME), using available RNA-seq data from three cohorts and the Cancer Genome Atlas database (**Table S11**). Therefore, cell networks of TME were constructed to depict the complex landscape of tumor-infiltrating lymphocytic cells, indicating that high HINS brought stronger active intercellular interactions in the responders (**Figure 4F-G**). Inspired by previous research [20], we constructed a gene expression profiling (GEP) score and a cytolytic (CYT) score, which were clustered in active immune responders. According to the immune score, improved levels of GEP- and CYT-related gene transcriptional expression were observed in patients with a high (vs. low) HINS (GEP: mean 3.26 vs. 2.97, CYT: mean 193.84 vs. 110.52; **Figure 5A**, **Table S12**). Furthermore, we performed a single-sample gene set enrichment analysis (ssGSEA) to study the associations between HINS distribution and specific immune molecules. Both the expression levels of HLA-I and HLA-II immune infiltration were elevated in high HINS responders (**Figure 5B**). Meanwhile, the infiltration of plasmacytoid dendritic cells (pDC) was amplified in the high HINS subgroup (positive correlation, 𝑅 = 0.124, 𝑃 = 1.5 × 10^-4^; **Figure S9B**). Additionally, the majority of immune checkpoint expression levels were compared across HINS subgroups to validate whether an improved response to ICB (identified by HINS) correlated with adaptive immune suppression. The results showed that the high HINS subgroup significantly enhanced the expression levels of evaluated checkpoint genes (PD-L1, PD1, CTLA4, LAG3, HAVCR2 and PDCD1LG2; **Figure S9A, C-G**, **Table S12**). This observation proved that adaptive immune activation had a higher priority than immune evasion in patients with ICB therapy.

**Figure 5.**
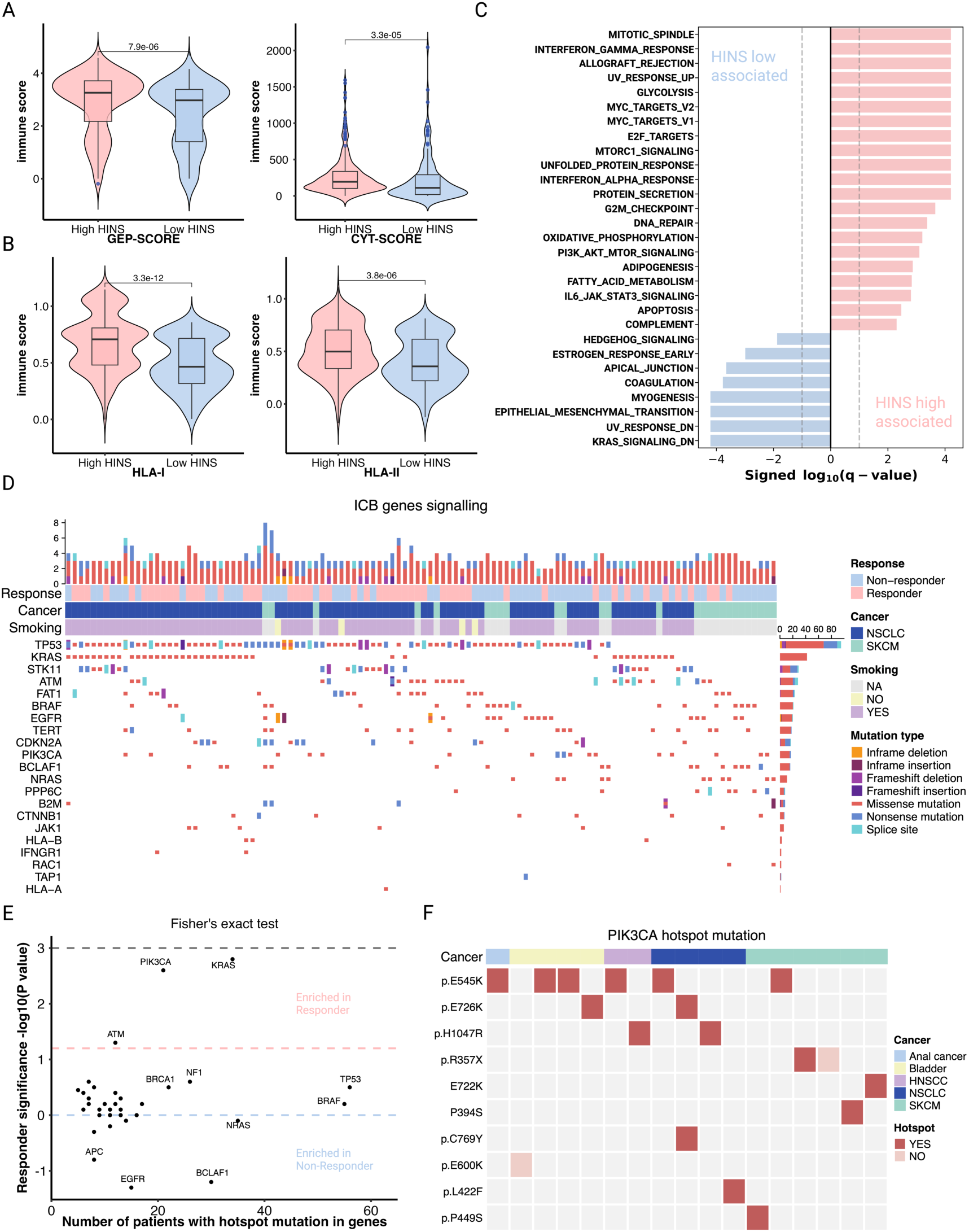
HINS is associated with multiple important transcriptional and genomic biomarkers. **A**. Violin plots of higher CYT-score and GEP-score in the high vs. low HINS subgroup (n = 455). **B**. Violin plots show significantly higher HLA I and II immune scores in the high (vs. low) HINS subgroup by single-simple gene set enrichment analysis. **C**. Hallmark gene set enrichment analysis of the response and resistance-associated pathways. **D**. CoMut plot of four constructed cohorts (comprised of NSCLC and SKCM) organized by response category. **E**. HINS-identified genes enriched in responder versus non-responder. **F**. Tile plots showing known hotspot and non-hotspot mutations in PIK3CA by distinct cancer types.

We performed gene set enrichment analysis based on Hallmark Gene Sets [48] to systematically infer differentially expressed gene pathways (𝑃 < 0.01, **Figure 5C**, **Table S13**). Top HINS-identified responder-associated pathways were enriched, including INTERFERON_GAMMA_RESPONSE, DNA_REPAIR and UV_RESPONSE_UP, which were reported as predictors of ICB response in previous studies. Pathways correlated with resistance were also obtained, including ESTROGEN_RESPONSE_EARLY and EPITHELIAL_MESENCHYMAL_TRANSITION (**Figure 5C**). Collectively, these results suggested that a high HINS may be correlated with immune transcriptional programs in patients with immunotherapy.

### HINS-identified mutational drivers associated with resistance

To facilitate a more comprehensive insight into the association between mutational drivers and HINS-identified response to ICB therapy, we analyzed the genomic-level features that could reveal further differences in this association. The prevalence of known drivers in common cancers (NSCLC and SKCM) was depicted (**Figure 5D**, **Table S14**). Meanwhile, we evaluated whether hotspot mutations in cancer driver genes were enriched in patients, as these events were more likely to influence tumor biology significantly (**Figure 5E**, **Table S15**). In all, clonal driver mutations in *PIK3CA*, *ATM*, and *KRAS* appeared to be highly favorable with respect to ICB response, while clonal driver alterations in *APC*, *EGFR*, *BCLAF1*, *NRAS* were unfavorable to patients with progressive disease. The tumor type distributions among *TP53*, *BRAF*, *BCLAF1* and *KRAS* were as expected, with TP53 displaying a pan tumor type distribution, *BCLAF1* and *BRAF* displaying a strong bias toward SKCM, and *KRAS* displaying a strong bias toward NSCLC.

Since driver events within a specific gene may arise in distinct functional domains and have different phenotypic impacts, we examined four representative genes (*PIK3CA*, *BCLAF1*, *TERT* and *KRAS*) for trends within specific cancer types (**Figure 5F**, **Figure S10**, **Table S16**). We observed that patients with clonal hotspot mutations in *PIK3CA* are associated with ICB response across almost all cancer types, especially for the missense mutation of *P.E545K*, which had a significant effect on cell proliferation [49]. As shown, *BCLAF1*, a factor that encoded the transcriptional repressor that regulates the type 1 interferon response [50], exhibited resistance to immunotherapy and was enriched in SKCM, NSCLC and bladder cancer. This observation was consistent with previous studies that mutations in the repressor gene *BCLAF1* were predictive of ICB non-responders [51]. Meanwhile, we found substantial non-hotspot mutations occurred within *TERT* in NSCLC, greatly increasing resistance in patients [28]. *KRAS* exhibited a similar trend with *PIK3CA*.

These results indicated that single-gene correlations with resistance to immunotherapy could provide abundant information to comprehensively clarify their relationships to tumors. Mechanistically, this correlation is underpinned by the ability of HINS to capture the intrinsic immunogenic quality of the neoepitopes derived from these hotspot mutations. Rather than just cataloging genomic alterations, HINS utilized the biophysical properties (e.g., HLA presentation and T cell cross-reactivity) of the resulting mutant peptides. For example, HINS identifies that the recurrent neoantigens generated by *PIK3CA* mutations possess a high-quality pattern conducive to T-cell recognition, whereas those from resistance genes like *BCLAF1* may lack such features. Thus, HINS effectively translates abstract genomic driver events into concrete quantitative immune signals, explaining their linkage to ICB survival benefits. Taken together, HINS-identified driver events may also provide the genomic insights of response or resistance to ICB therapy.

## Discussion and conclusion

The complexity of infiltrated tumor microenvironment and immunopeptidome diversity emphasize the necessity for comprehensive models of immunotherapy response prediction. Modeling the quality of immunogenic neoantigens that differentially bind to HLA and activate T cells is crucial to measuring the strength of the immune response. Herein, we introduced a predictor of immune checkpoint blockade response, namely HLA-I-mediated immunogenic neoantigen score, using predicted neoantigen quality and evolutionary divergence of HLA to integrate the aspects with immune activation and evasion. This method used a deep attention model to merge multi-modal features, such as ‘non-self’ recognition, presentation of HLA, and cross-reactivity of T cells, along with establishing an interpretable score algorithm that is predictive of stratification of patients with ICB therapy.

Previous studies have demonstrated that antigen presentation is compromised because of the LOH at HLA loci, increased divergence of HLA-I allele across varied HLA-I subtypes, or loss of B2M may allow immune escape in certain types. Our results provide a rationale for merging HLA-I allele divergence and multi-modal features. Specifically, HINS yields an overall accuracy of AUC=0.853, outperforming existing predictors and capturing more than 75% true responders, greatly reducing the false discovery rate. Meanwhile, HINS exhibited compelling predictive performance across different cancer types (e.g., NSCLC, SKCM, and HNSCC) and a declined HR compared with ICB-associated biomarkers (TMB, HED). Participants with high HINS were more likely to generate high-quality immunogenic neoantigens that can strongly trigger the anti-tumor immune response.

Further Kaplan-Meier analysis in TMB and HLA-LOH revealed three viewpoints: 1) HINS successfully identified responders though in the low TMB population, 2) HINS improved the prediction performance with HLA-LOH status, and 3) HINS significantly correlated with signal of actual TCR immune response and enhance the predictive power of ICB response. Thus, the combination of HINS with TMB, HLA-LOH or TCR diversity proved the clear fact that HINS is a powerful predictor with high resolution to stratify patients treated by immunotherapy. Transcriptomic analysis in our constructed cohorts was notable for the identification of the relationship between HINS and the immune transcriptional programs in the tumor microenvironment. A high HINS was correlated with elevated levels of immune cells, immune checkpoints, and differentially expressed gene pathways. The genomic-level features were also analyzed to distinguish important mutational driver genes across different cancer types. Of the top 21 HINS-identified genes, four (*PIK3CA*, *BCLAF1*, *TERT* and *KRAS*) were representatively associated with response or resistance to immunotherapy. Taken together, HINS could be a reliable tool for enhancing decision-making practices in ICB therapy to maximize patient survival and provide transcriptional and genomic insights into underlying biological tumor mechanisms.

There are a few limitations in this study. Firstly, we have a retrospective nature of method design. Although we have comprehensively integrated the multi-modal biological features in immune response, the heterogeneity and complexity of tumors is still intractable to fully characterize. Thus, more prospective studies need to be carried out in the future. Secondly, we only enrolled patients across six cancer types, with NSCLC and SKCM accounting for most samples. Although the current results supporting HINS is strongest for these tumors, further validation in under-represented cancer types is still needed. Furthermore, the utilization of multiple cohorts inevitably introduces potential batch effects arising from heterogeneous sequencing platforms and clinical regimens. To mitigate technical variations, we employed a unified, standard pipeline to process raw WES data across all cohorts, ensuring consistent mutation calling. Distinct from transcriptomic signatures susceptible to quantitative sequencing noise, HINS relies on intrinsic physicochemical properties of neoantigen sequences and HLA evolutionary divergence. Nevertheless, residual batch effects cannot be completely excluded, and future prospective datasets with fully standardized pipelines will further validate its robustness. Thirdly, the biomarker PD-L1 expression is observed to improve the predictive power of immunotherapy response but is strictly limited by the sample size. Whereas HINS was available for the entire WES cohort. Therefore, further evaluation needs to be conducted to confirm its importance in predicting ICB outcomes with extensive cohorts [52]. Fourthly, the definition of neoantigen immunogenicity in our validation cohorts relied on a binding affinity threshold rather than experimentally verified T-cell responses. This may potentially missing intricate biochemical patterns specific to TCR recognition. Future iterations of HINS will benefit significantly from training on large-scale datasets with ground-truth neoantigen immunogenicity. Additionally, the genomic and transcriptional included in this study solely at the bulk-RNA level. and thus, we could not further discuss the recognition and evasion mechanism of different cell types within TME. While we validated the synergy between HINS and TCR repertoire diversity, our dependence on bulk RNA-seq data still limits the ability to decouple tumor-reactive clonotypes from bystander T cells. Because the shape of single-cell transcriptomics data [53] is hard to unify with our data. Consequently, a pivotal direction for future research lies in integrating HINS with matched single-cell TCR sequencing. Lastly, class II HLA neoantigens are not analyzed in this study and should be explored in the future to provide the whole landscape of HLA-mediated anti-tumor responses.

In summary, through integrated analysis of HLA evolutionary divergence and immunogenic neoantigen quality, we introduce an interpretable deep learning-based model to predict patients’ objective response following immunotherapy. HINS merges multi-modal features and correlates with crucial biomarkers, which exhibit great potential to facilitate patient stratification. In full awareness of this perception, we expect to see our study and other analogous studies could inspire more formal and prospective findings in personalized precision therapy with broader genomic assays.

## Code availability

HINS is an open access tool that available at https://github.com/TomasYang001/HINS.

## Acknowledgements

The authors acknowledge support from Natural Science Foundation of Heilongjiang Province of China (ZD2024F001), National Natural Science Foundation of China (Nos. 62225109, 62450122, 62372135, 62202092), and the King Abdullah University of Science and Technology (KAUST) Office of Research Administration (ORA) under Award No REI/1/5234-01-01, REI/1/5414-01-01, REI/1/5289-01-01, REI/1/5404-01-01, REI/1/5992-01-01, URF/1/4663-01-01, Center of Excellence for Smart Health (KCSH), under award number 5932, Center of Excellence on Generative AI, under award number 5940.

---

**Figure S1.**
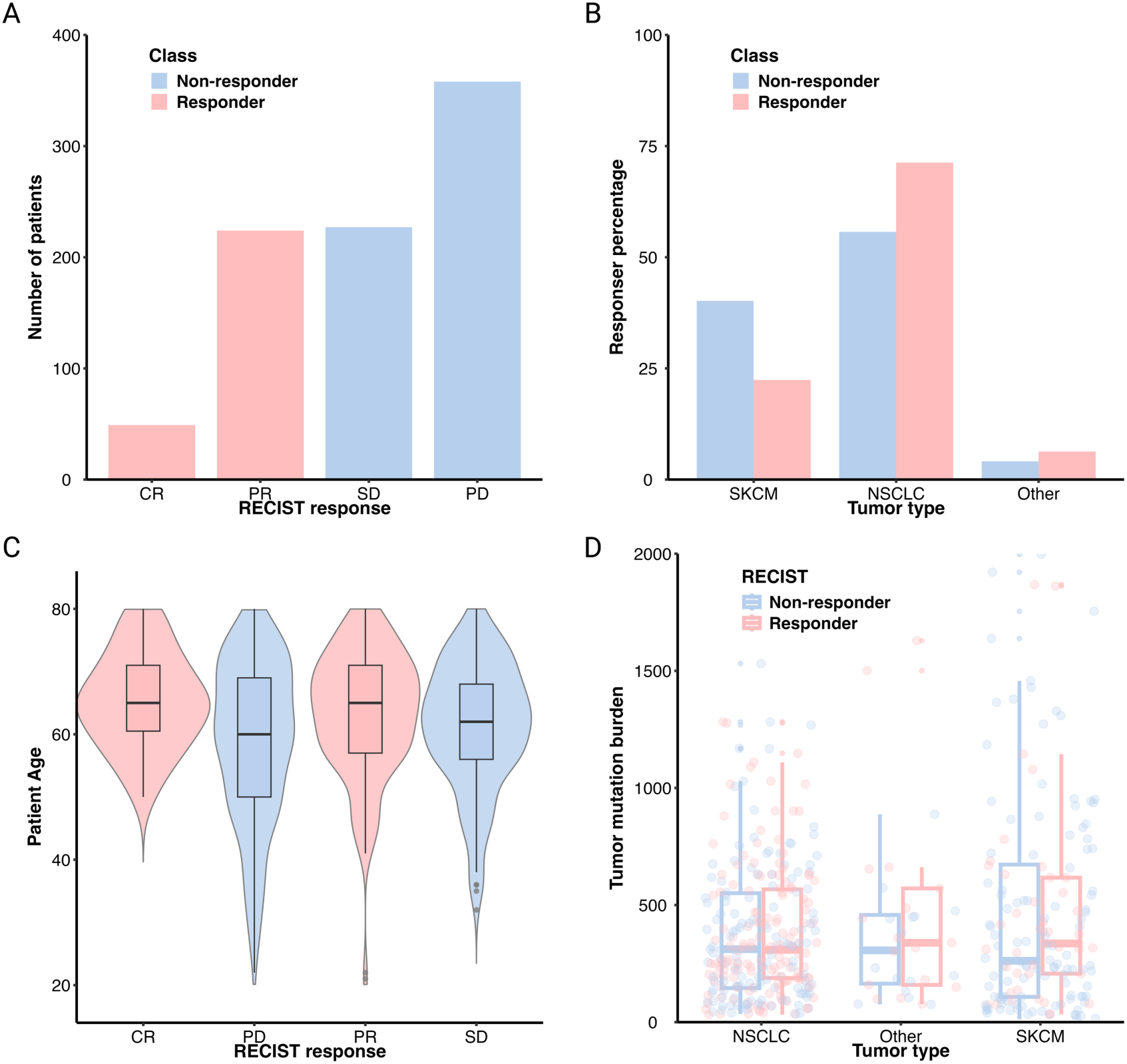
Clinical characteristics of curated patients with immune checkpoint blockade therapy. **A**. Number of patients from the aggregated nine cohorts in each RECIST response group. **B**. Proportion of tumor types in immune checkpoint blockade responders and non-responders. **C**. Patient ages for different RECIST response groups. **D**. TMB for responders and non-responders to immunotherapy by tumor type. Statistical significance was tested using two-tailed Welch’s t-tests.

**Figure S2.**
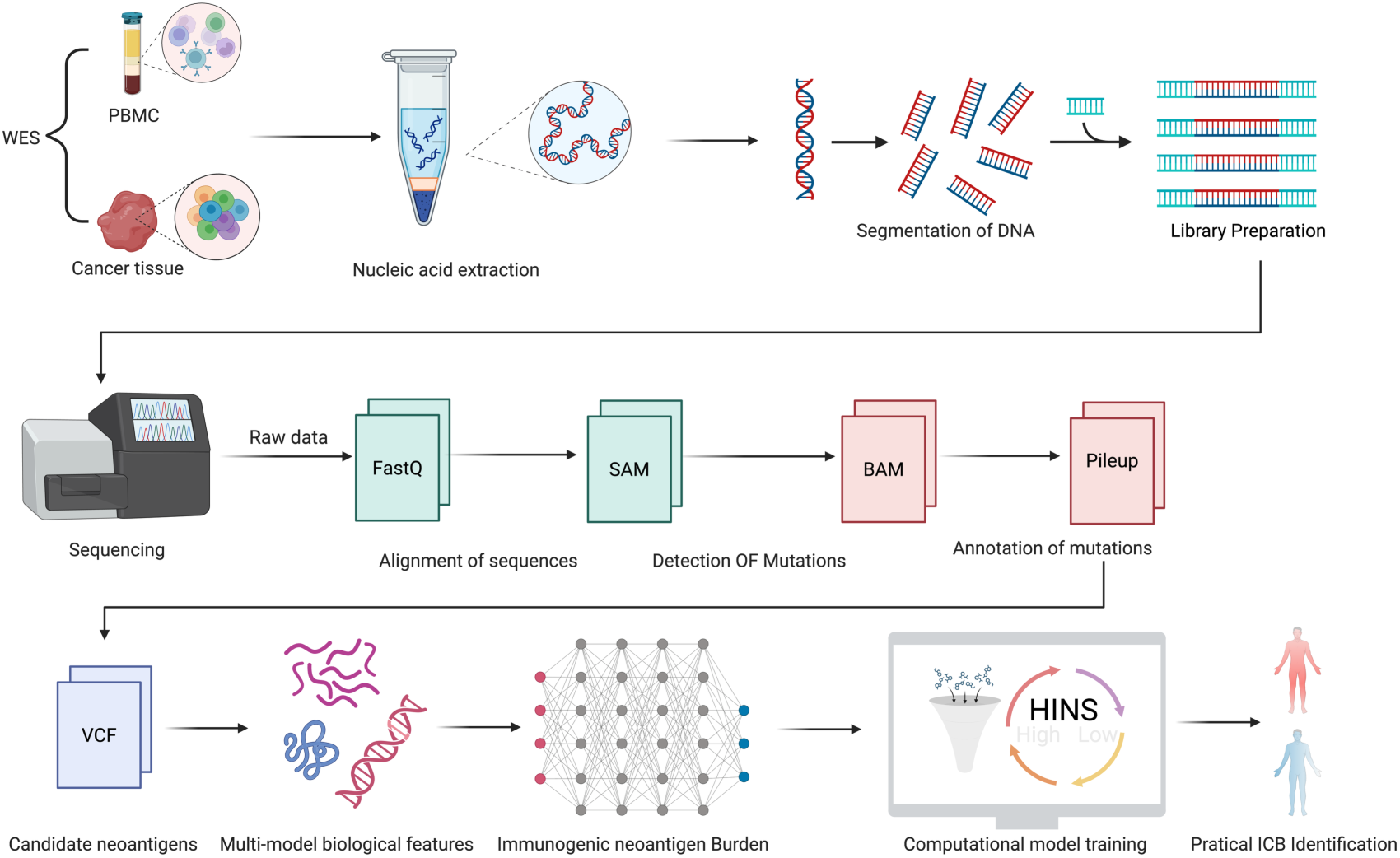
Overview of the pipeline used to process the whole-exome sequencing data of ICB cohorts.

**Figure S3.**
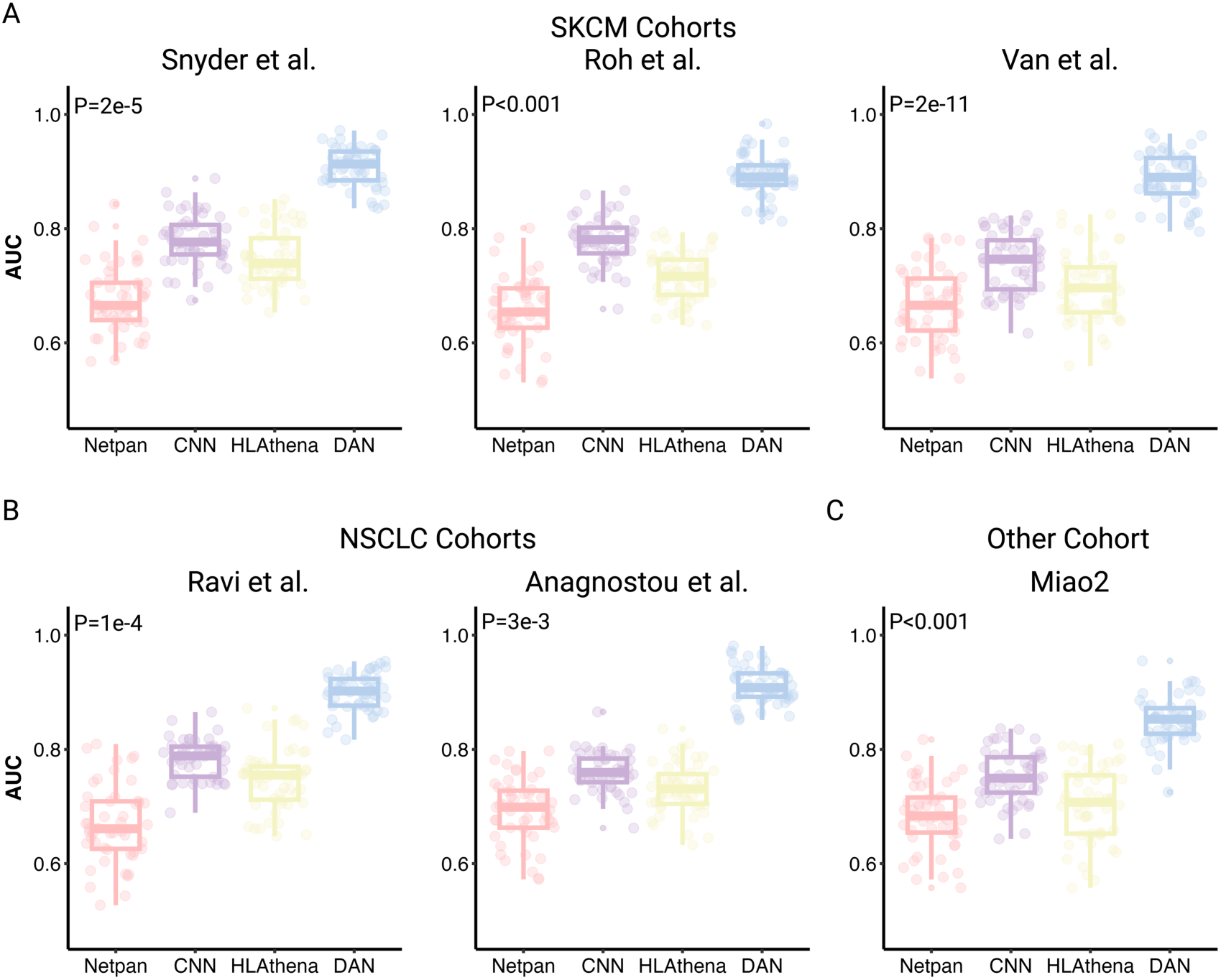
Box plots of AUC results for tumor-specific neoantigen prediction across multiple cohorts. **A**. Comparison of our proposed deep attention networks to other state-of-the-art tools in SKCM cancer, i.e., NetMHCpan-4.1 (represented as Netpan), CNN, and HLAthena. **B**. Comparison in NSCLC cancer. **C**. Comparison in pan-cancer in Miao et al.’s cohort, including bladder cancer, head and neck squamous cell carcinoma, anal cancer, and sarcoma.

**Figure S4.**
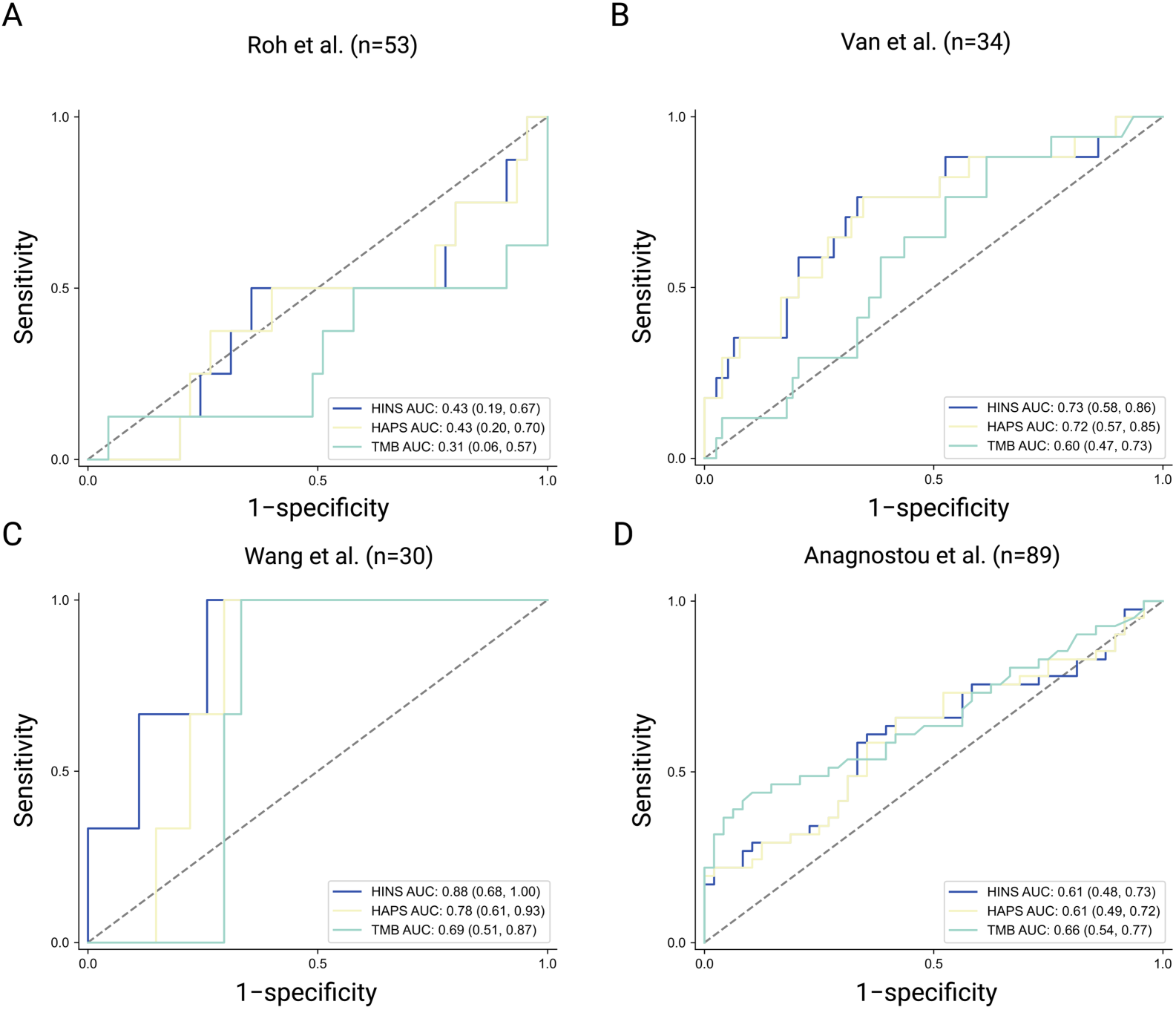
Predictive performance of HINS, HAPS and the TMB biomarker for ICB response. **A**. AUC plot of HINS, HAPS and TMB in predicting ICB response in SKCM cancer. Roh et al.’s cohort, n=53. **B**. AUC plot of HINS, HAPS and TMB in predicting ICB response in SKCM cancer. Van et al.’s cohort, n=34. **C**. AUC plot of HINS, HAPS and TMB in predicting ICB response in NSCLC cancer. Wang et al.’s cohort, n=30. **D**. AUC plot of HINS, HAPS and TMB in predicting ICB response in NSCLC cancer. Anagnostou et al.’s cohort, n=89.

**Figure S5.**
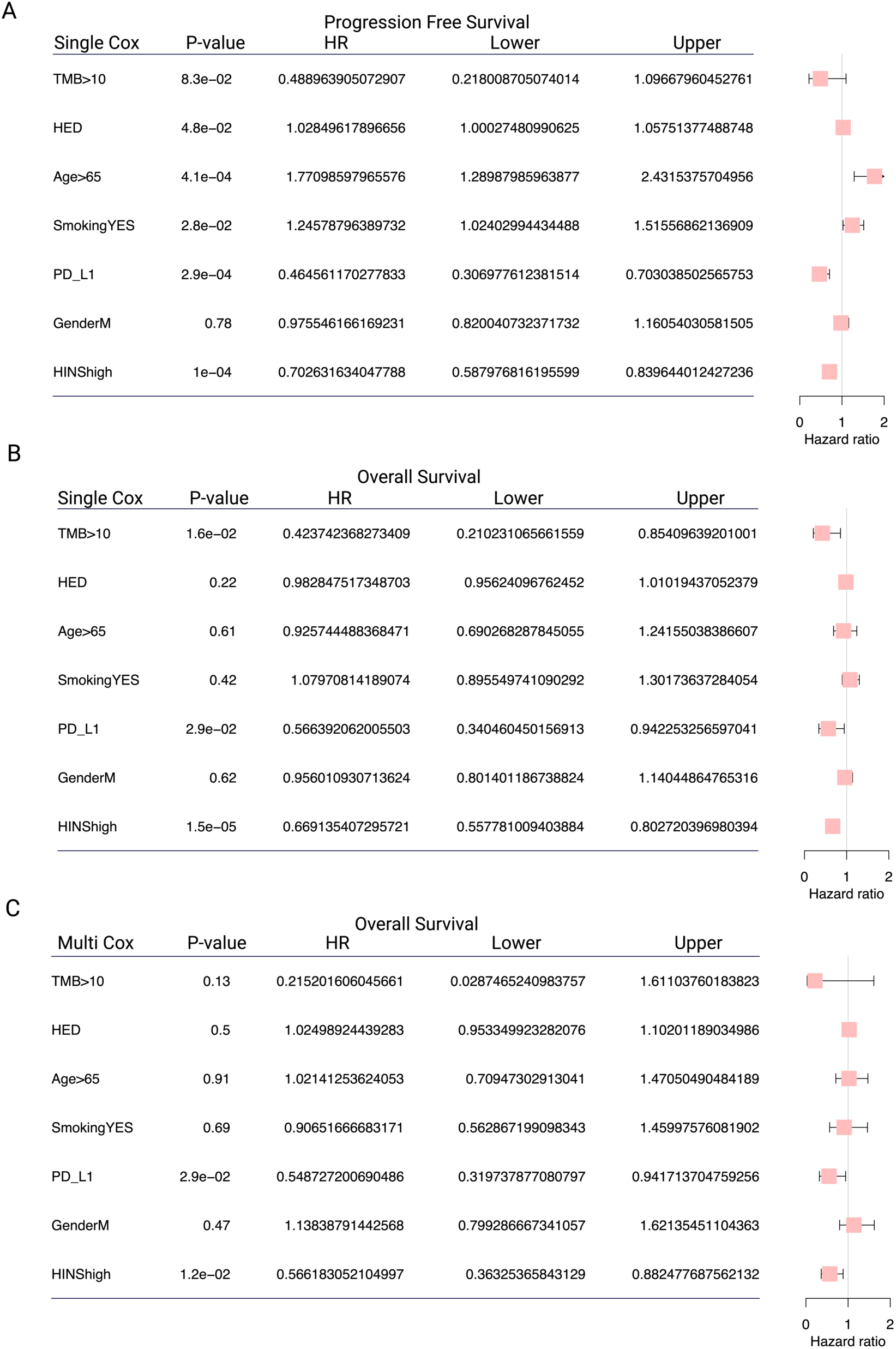
Univariate and multivariate Cox proportional hazards regression analysis. **A**. Significantly prolonged PFS in the high HINS subgroup by univariate Cox analysis. **B-C**. Greatly improved OS in the high HINS subgroup by univariate (B) and multivariate (C) Cox analysis.

**Figure S6.**
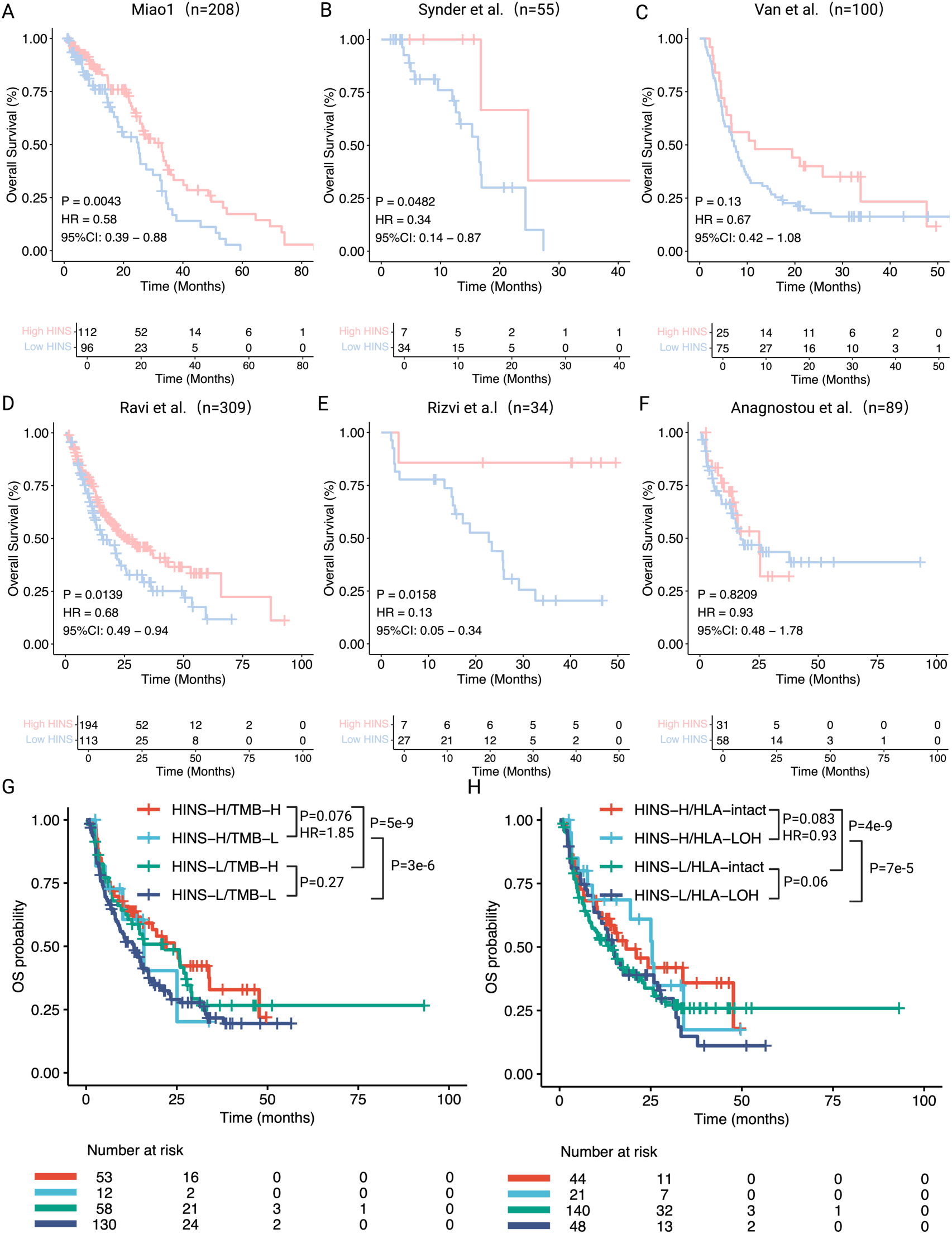
HINS predicts patient OS outcomes following ICB therapy with powerful stratification. **A-F**. Kaplan–Meier (K-M) analysis shows significant associations between high HINS and improved patient OS after ICB therapy across distinct cohorts. Miao et al.’s SKCM cohort, n=208 (A); Synder et al.’s SKCM cohort, n=55 (B); Van et al.’s SKCM cohort, n=100 (C); Ravi et al.’s NSCLC cohort, n=309 (D); Rizvi et al.’s NSCLC cohort, n=34 (E); Anagnostou et al.’s NSCLC cohort, n=89 (F). **G-H**. K-M analysis of OS when combining HINS and TMB (G) or HLA-LOH (H). The P-value was determined by univariable Cox proportional hazards regression.

**Figure S7.**
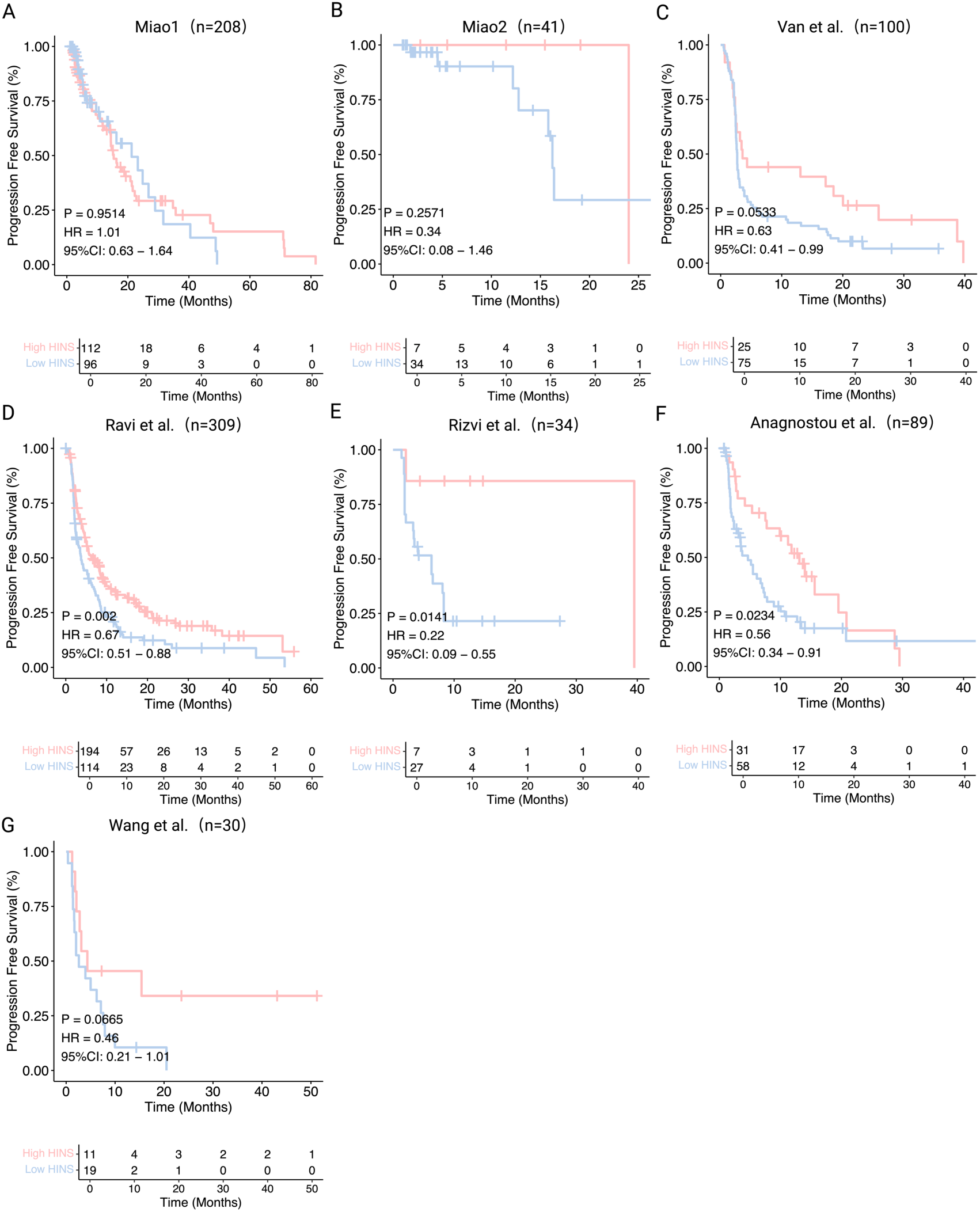
HINS predicts patient PFS outcomes following immunotherapy with powerful stratification. **A-G**. Kaplan–Meier (K-M) analysis shows significant associations between high HINS and improved patient OS after ICB therapy across multiple cancers. Miao et al.’s SKCM cohort, n=208 (A); Miao et al.’s cohort including multiple cancers, n=41 (B); Van et al.’s SKCM cohort, n=100 (C); Ravi et al.’s NSCLC cohort, n=309; Rizvi et al.’s NSCLC cohort, n=34 (E); Anagnostou et al.’s NSCLC cohort, n=89 (F); Wang et al.’s NSCLC cohort, n=30 (G).

**Figure S8.**
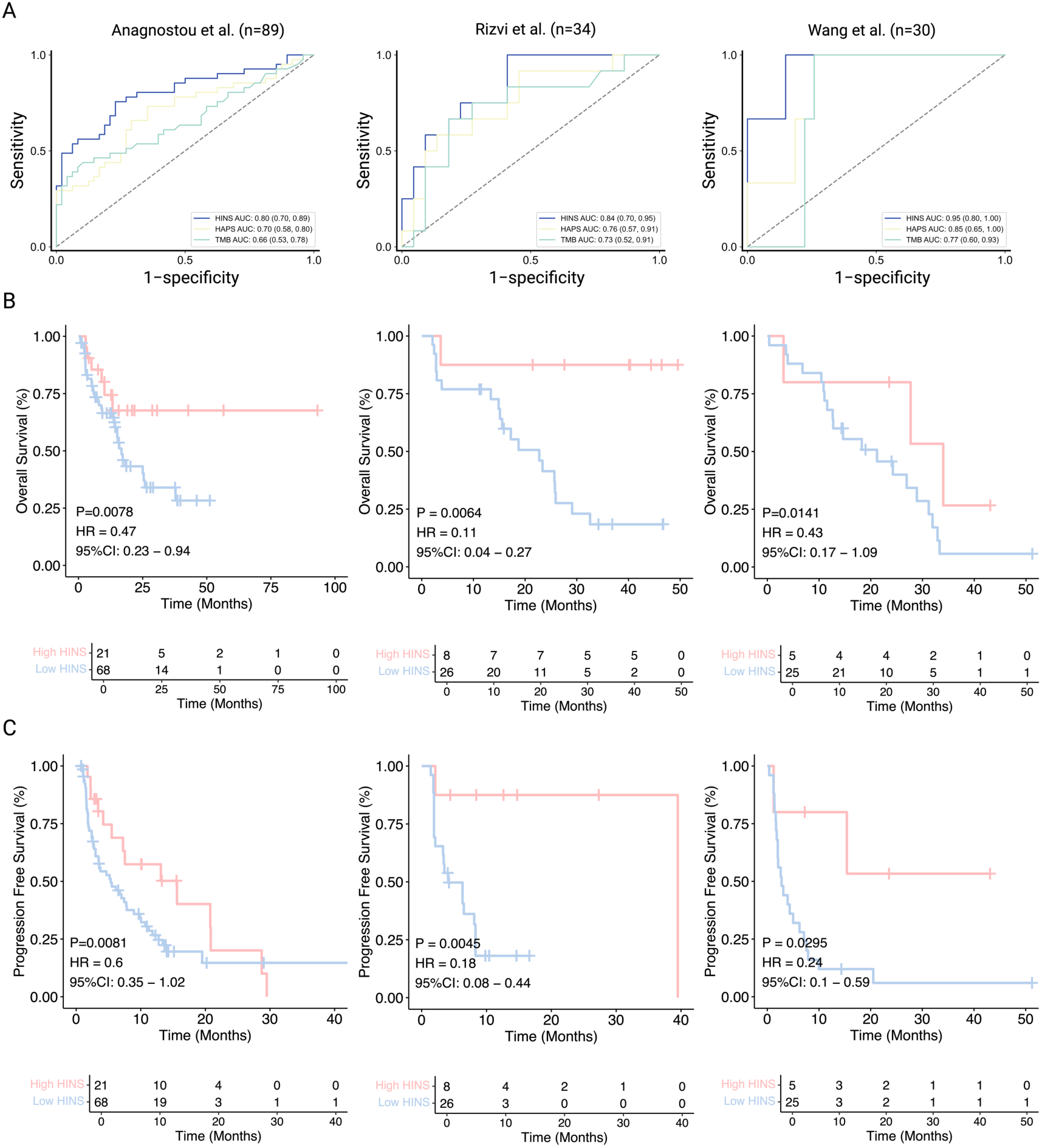
Performance of the NSCLC-specific HINS model in three independent NSCLC cohorts. **A**. ROC curves of the NSCLC-specific HINS score (blue) for predicting ICB response in the Anagnostou et al. (n = 89), Rizvi et al. (n = 34) and Wang et al.’s (n = 30) NSCLC cohorts, compared with HAPS (yellow) and the TMB biomarker (green). **B**. Kaplan-Meier OS curves stratified by high versus low HINS (cutoff = 18.5) in the three NSCLC cohorts; corresponding log-rank P values, hazard ratios (HRs) and 95% confidence intervals (CIs) are indicated. **C**. Kaplan-Meier PFS curves for the same NSCLC cohorts, further demonstrating that higher HINS is associated with improved clinical outcomes under immune checkpoint blockade.

**Figure S9.**
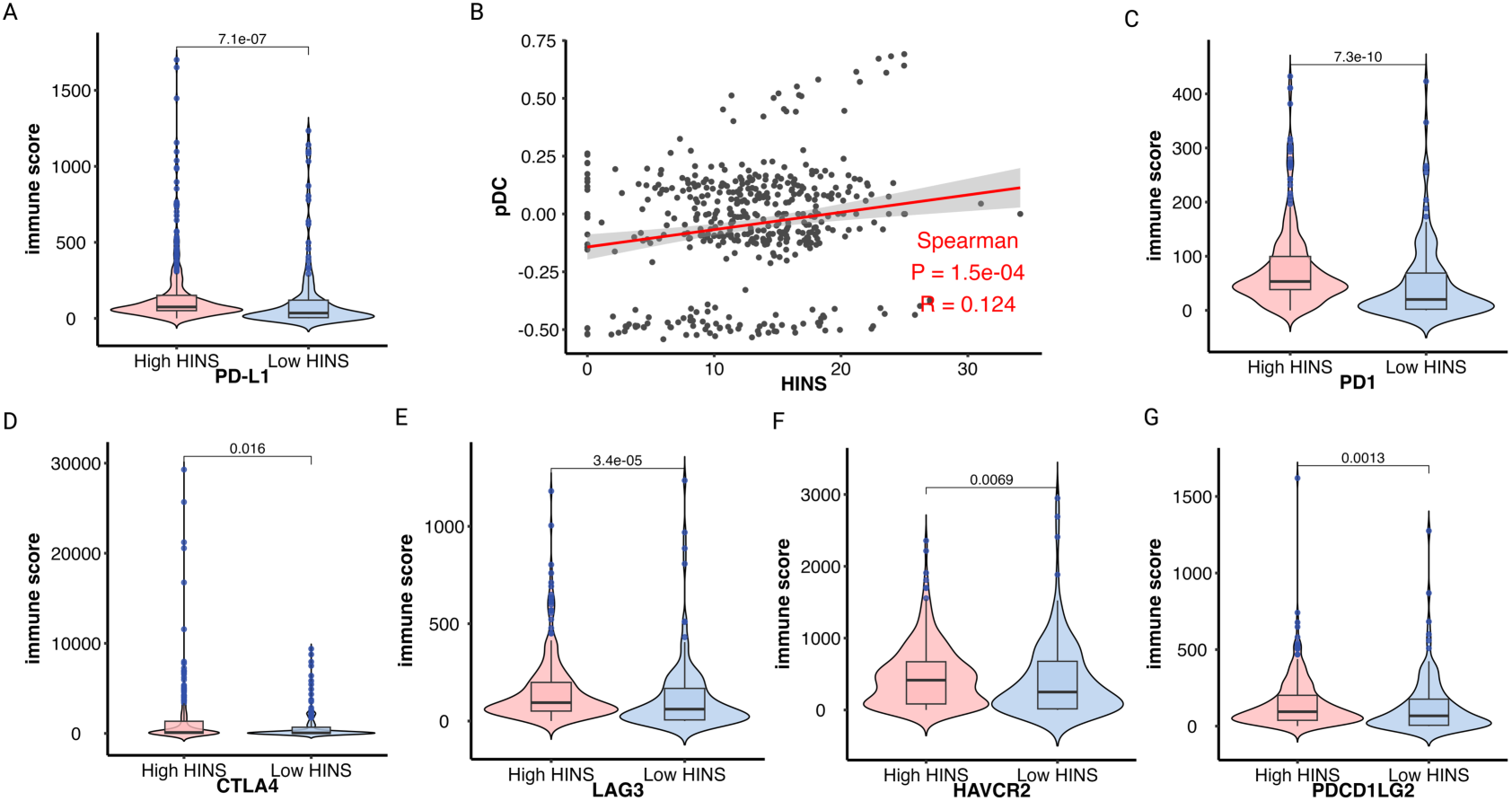
Immune microenvironment differences between HINS and checkpoints. **A**. Significantly higher expression levels of PD-L1 in the high HIN subgroup. **B**. A positive association between plasmacytoid dendritic cell infiltration level and HINS by ssGSEA (two-tailed). Shaded area, 95% CI for the association. **C-G**. The high HINS subgroup had significantly higher expression levels of immune checkpoint genes (PD1, CTLA-4, LAG3, HAVCR2, PDCD1LG2).

**Figure S10.**
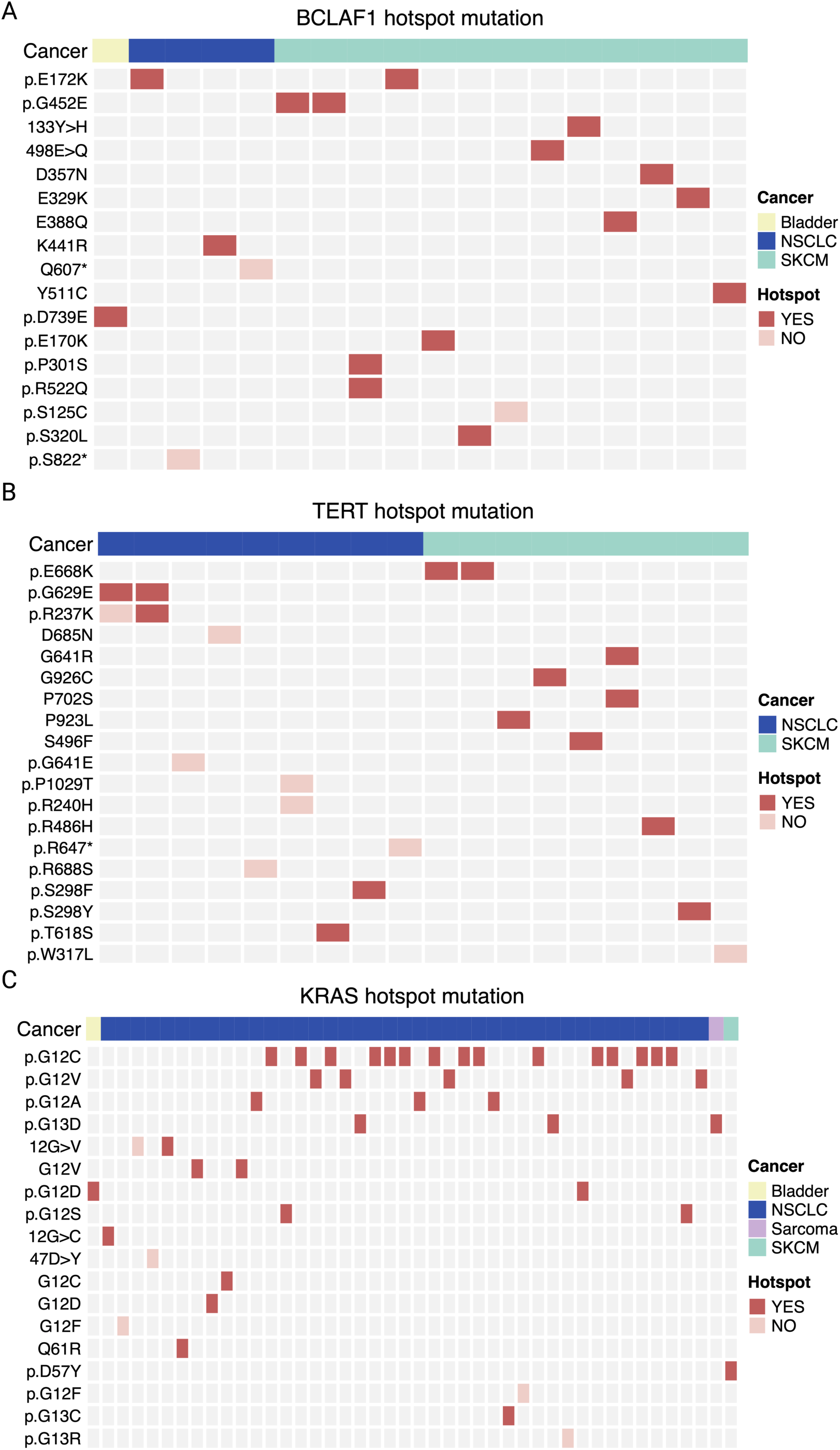
Tile plots showing known hotspot and non-hotspot mutations in HINS-identified genes. **A.** BCLAF1 hotspot mutation. **B.** TERT hotspot mutation. **C.** KRAS hotspot mutation.

