## Supplemental Table 1-16 for "HINS: HLA-mediated immunogenic neoantigen score for robust prediction of immune checkpoint blockade response": Table S2.docx

**Table S2 The detailed parameter settings of deep attention networks**

| Hyper-parameters | Values |
| --- | --- |
| Peptide length $\mathcal{l}$ | 17 |
| Embedding dimension $\nu$ | 20 |
| MTE layers | [2, 3, 4] |
| MTE hidden length | 1300 |
| MTE attention | [2, 4, 10] |
| MTE dropout | 0.1 |
| MIL peptide input | [17, 1300] |
| MIL HLA input | [35, 1300] |
| MIL interaction layers | 1 |
| MIL pooling layers | 1 |
| Fully neural networks | 4 |
| $\varphi$ | 0.224 |

*Note*: $\varphi$ indicates the relative weight between the terms of presentation and cross-reactivity. MTE is the molecule transformer encoder block. MIL is the mutual interaction learning block.
