## Supplemental Table 1-16 for "HINS: HLA-mediated immunogenic neoantigen score for robust prediction of immune checkpoint blockade response": Table S5 .docx

**Table S5 Comparison of three variants among SKCM and NSCLC cancer regarding AUC, AUPR, PPVn and F1-score values**

|  | Method | AUC | AUPR | PPV | F1-score |
| --- | --- | --- | --- | --- | --- |
| SKCM | BLOSUM62 | 0.851(0.004) | 0.413(0.012) | 0.502(0.009) | 0.612(0.006) |
|  | ESM2 | 0.862(0.004) | 0.427(0.010) | 0.524(0.007) | 0.617(0.005) |
|  | DAN | **0.873(0.003)** | **0.437(0.009)** | **0.532(0.007)** | **0.633(0.005)** |
| NSCLC | BLOSUM62 | 0.827(0.003) | 0.413(0.013) | 0.502(0.007) | 0.611(0.005) |
|  | ESM2 | 0.834(0.003) | 0.416(0.012) | 0.506(0.007) | 0.619(0.004) |
|  | DAN | **0.842(0.002)** | **0.425(0.011)** | **0.513(0.006)** | **0.627(0.004)** |

*Note*: Best result is bolded. BLOSUM62 indicates the input of DAN model is embedded by BLOSUM62. ESM2 indicates the input of DAN model is embedded by ESM2.
